# Multilayer regulation underlies the functional precision and evolvability of the olfactory system

**DOI:** 10.1101/2025.01.16.632932

**Authors:** Jérôme Mermet, Steeve Cruchet, Asfa Sabrin Borbora, Daehan Lee, Phing Chian Chai, Andre Jang, Karen Menuz, Richard Benton

## Abstract

Sensory neurons must be reproducibly specified to permit accurate neural representation of external signals but also able to change during evolution. We studied this paradox in the *Drosophila* olfactory system by establishing a single-cell transcriptomic atlas of all developing antennal sensory lineages, including latent neural populations that normally undergo programmed cell death (PCD). This atlas reveals that transcriptional control is robust, but imperfect, in defining selective sensory receptor expression. A second layer of precision is afforded by the intersection of expression of functionally-interacting receptor subunits. A third layer is defined by stereotyped PCD patterning, which masks promiscuous receptor expression in neurons fated to die and removes “empty” neurons lacking receptors. Like receptor choice, PCD is under lineage-specific transcriptional control; promiscuity in this regulation leads to previously-unappreciated heterogeneity in neuronal numbers. Thus functional precision in the mature olfactory system belies developmental noise that might facilitate the evolution of sensory pathways.

## Introduction

Sensory systems mediate detection of the environment and provide the brain with a spatio-temporal code that enables recognition, interpretation and appropriate behavioral responses to a stimulus. However, the external world of stimuli changes as species colonize new ecological niches. Thus, sensory systems must also be capable of change over evolutionary timescales.

This paradox of functional precision but evolutionary flexibility is particularly notable in the olfactory system of *Drosophila melanogaster*. Intensive anatomical, molecular and functional analyses of the major olfactory organ, the third antennal segment (hereafter, antenna), have defined a highly stereotyped organization in which ∼1200 neurons are categorized into nearly 50 distinct classes of olfactory sensory neurons (OSNs), as well as several types of hygrosensory and thermosensory neurons (Benton, 2022; Benton et al., 2025; Couto et al., 2005; Li et al., 2022; Li et al., 2020; McLaughlin et al., 2021; Schlegel et al., 2021; Vosshall and Stocker, 2007). Each class of olfactory neuron is characterized by the expression of a specific “tuning” receptor (belonging to the Odorant receptor (Or), Ionotropic receptor (Ir) or so-called “Gustatory” receptor (Gr) families), which defines chemical specificity, together with one or more broadly-expressed “co-receptors” (Orco for Ors; Ir8a, Ir25a, Ir76b for Irs). A few classes of neurons express more than one tuning receptor, typically encoded by tandemly-arranged gene duplicates. Antennal sensory neurons are grouped in combinations of 1-4 neurons underlying sensory hairs (sensilla) of several distinct morphological classes: antennal basiconic (ab), trichoid (at), intermediate (ai) and coeloconic (ac) (Figure 1A). The ciliated dendrites of OSNs are housed in the sensillar hair, while the axons project to the antennal lobe in the brain, where they innervate a glomerulus unique to each type of neuron.

**Figure 1.**
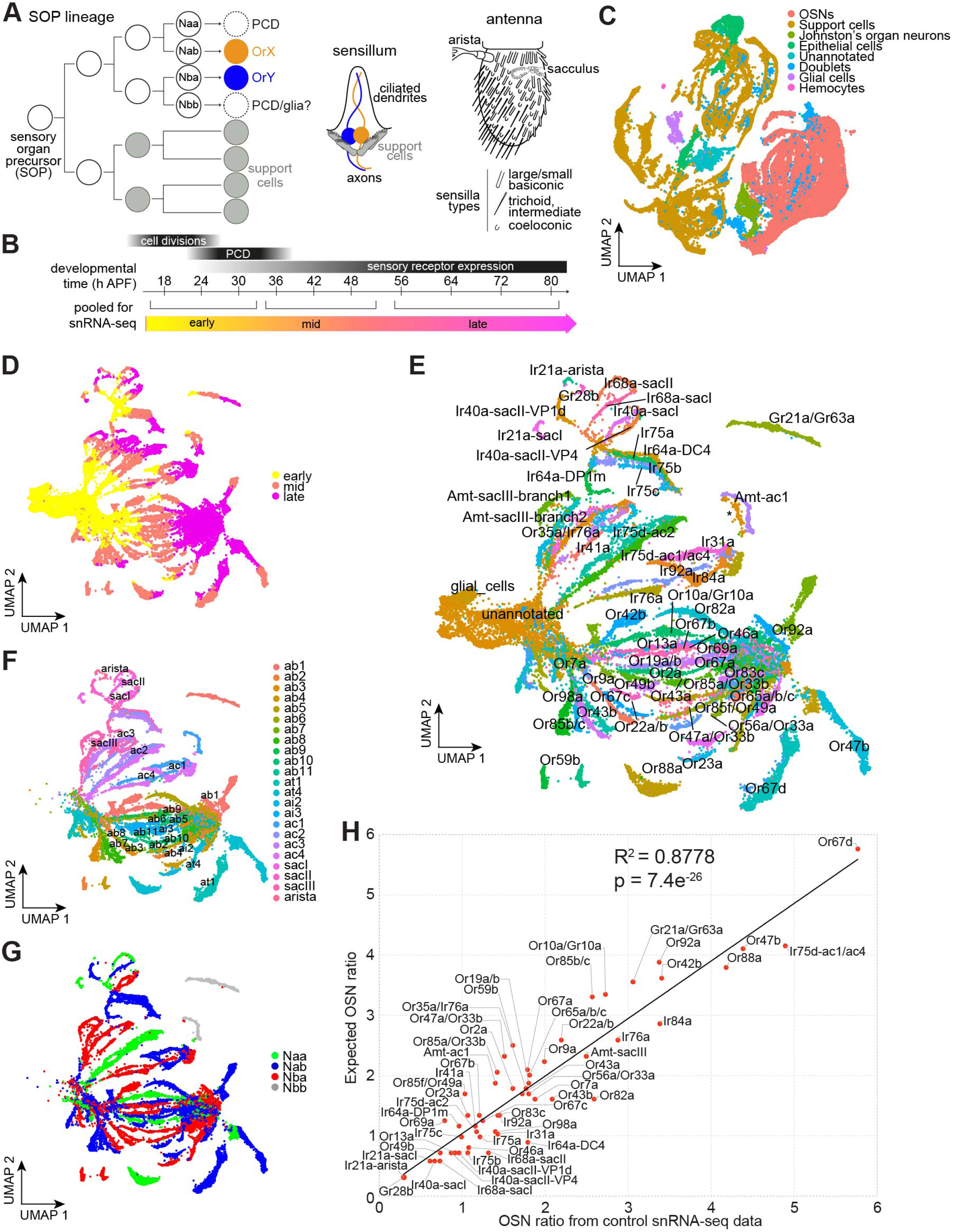
A developmental atlas of antennal sensory neurons. **(A)** Schematic of the development and anatomy of the *D. melanogaster* peripheral olfactory system. **(B)** Schematic of experimental design: antennal imaginal discs/antennae were dissected at time points every 6-8 h from 18-80 h after puparium formation (APF) in control (*peb-Gal4/+;;UAS-unc84:GFP/+*) and PCD-blocked (*peb-Gal4/+;UAS-p35/+;UAS-unc84:GFP/+*) conditions. Samples were pooled into three temporal phases (early, mid, late) prior FACS sorting and 10x sequencing. Tapered black bars indicate the timing of the main developmental processes. **(C)** UMAP of all cell types in the developmental atlas, integrating control and PCD-blocked datasets. **(D)** UMAP of all olfactory sensory neurons in (C) in the atlas – integrating control and PCD-blocked datasets – colored by developmental phases. **(E)** UMAP of annotated neuronal lineages, integrating control and PCD-blocked datasets. Unannotated neurons could not be assigned to any lineage; almost all of these are from early developmental stages, but one cluster (specific to the PCD-blocked dataset) was detected in late developmental stages (asterisk, close to the Amt-ac1 lineage). Some *repo* expressing glial cells were also detected (see Figure S3A-B). **(F)** UMAP of the cells in (E) – masking unannotated neurons and glial cells – colored by sensillar class (ab, antennal basiconic; at, antennal trichoid; ai, antennal intermediate; ac, antennal coeloconic; sac, sacculus). **(G)** UMAP of the cells in (F) – now also masking cells from the PCD-blocked dataset and dying and aristal lineages from the control dataset – colored by neuron precursor type (aristal lineages could not be confidently assigned to any type). **(H)** Scatter plot of the relative abundance of each neuronal population in the developmental atlas (control dataset only) with their relative abundance as quantified *in situ* (Benton et al., 2025).

The stereotypy is derived from apparently hard-wired developmental mechanisms. Each sensillum develops from a single sensory organ precursor (SOP) cell that is specified in the antennal imaginal disk in the larva (Figure 1A). The canonical view is that an SOP gives rise to a short, fixed lineage of asymmetric cell divisions that produces eight terminal cells with distinct molecular identity (Chai et al., 2019; Endo et al., 2007; Endo et al., 2011). Four of these eight cells become support cells (which have functions in sensillum construction and secretion of perireceptor proteins (Schmidt and Benton, 2020)), while the other four – termed Naa, Nab, Nba or Nbb – can potentially give rise to OSNs (Chai et al., 2019; Endo et al., 2011). Although two sensillum types do contain 4 OSNs, all other sensilla house fewer neurons. This is thought to be due to programmed cell death (PCD) of precursor cells during the pupal stage (Chai et al., 2019; Endo et al., 2007; Endo et al., 2011); in many ac lineages the Nbb precursor is thought to differentiate as a glial cell (Endo et al., 2007; Rodrigues and Hummel, 2008; Sen et al., 2005). Abundant evidence supports the contribution of OSN-specific gene regulatory networks in defining the fate of surviving neurons, notably in the precise transcriptional activation (or inhibition) of receptor genes (Barish and Volkan, 2015; Jafari et al., 2012; Mika and Benton, 2021; Mika et al., 2021). Such deterministic transcriptional codes are thought to be central to the functional stereotypy of the olfactory system.

Comparison of the *D. melanogaster* olfactory system with other insects, however, reveals remarkable evolvability, with changes in receptor function, receptor expression and OSN number, often linked to adaptation of species to new ecological niches (Hansson and Stensmyr, 2011; Ramdya and Benton, 2010; Zhao and McBride, 2020). For example, in *Drosophila sechellia*, an extreme specialist on noni fruit, olfactory channels detecting the host fruit exhibit altered receptor tuning and expanded OSN populations (Auer et al., 2020; Prieto-Godino et al., 2017; Takagi et al., 2024).

The generation of new receptors through tandem gene duplication and functional diversification through sequence changes are conceptually straightforward processes that are well-documented (Croset et al., 2010; Mika et al., 2021; Robertson et al., 2003). By contrast, how novel cell types within a sensillum might emerge is much less well-understood. One clue came from the demonstration that blocking PCD is sufficient to result in the formation of functional OSNs (Prieto-Godino et al., 2020), implying a latent potential of OSNs fated to die in evolving as new cell types. Deeper understanding of this potential is precluded by our almost complete lack of knowledge of how PCD is patterned in the developing OSN lineages, and the molecular properties of individual neurons fated to die.

In this work, we generate a high-resolution, developmental atlas of the antennal neuronal lineages, encompassing those that become functional neurons as well as those that undergo PCD. We use this to define the first molecular determinants specifying PCD of OSNs. Notably, we also discovered previously overlooked heterogeneity in the patterning of receptor expression and PCD in the olfactory system, which suggests how it might adapt during evolution.

## Results

### Generation of an atlas of developing and dying antennal sensory neurons

To generate a high-resolution, comprehensive, spatio-temporal atlas of the antennal OSN lineages, we labelled these cells by using *pebbled-Gal4* (*peb-Gal4*), which is expressed in all neural lineages (as well as many non-neuronal cells) from before 18 h after puparium formation (APF) (Sweeney et al., 2007), to drive a nuclear-GFP reporter (*UAS-unc84:GFP*) (Henry et al., 2012). In parallel, to characterize the developmental potential of neurons that are ultimately lost due to PCD during ∼22-32 APF (Chai et al., 2019; Endo et al., 2007; Prieto-Godino et al., 2020; Sen et al., 2004), we blocked OSN death by using *peb-Gal4* to also drive *UAS-p35*, encoding the baculoviral P35 caspase inhibitor (Prieto-Godino et al., 2020). For both control and PCD-blocked genotypes, we dissected antennal tissues from animals sampled every 6-8 h from 18-80 h APF, spanning the vast majority of their development (Figure 1B). Antennae were pooled into “early” (18-30 h APF), “mid” (36-48 h APF) and “late” (56-80 h APF) developmental stages prior to FACS isolation of GFP-positive nuclei (Figure 1B). Using the 10X Genomics Chromium platform, we sequenced the transcriptomes of ∼54k and ∼32k nuclei from control and PCD-blocked antennae, respectively, detecting on average ∼1000 genes/nucleus (Figure S1A).

Unless mentioned otherwise (see Methods and legends), for all downstream analyses we integrated control and PCD-blocked datasets, assuming that the vast majority of cells would form equivalent clusters in these datasets, and that a smaller number of “undead cells” would potentially form clusters unique to the PCD-blocked dataset. To broadly catalog antennal cell types, we used marker genes extracted from the Fly Cell Atlas (Li et al., 2022) (see Methods) (Figure S1B-C). Sensory neurons – excluding Johnston’s organ mechanosensory neurons – represent ∼39% of cells (∼21,000) in the control dataset and ∼43% (∼14,000 cells) in the PCD-blocked dataset (Figure S1D), consistent with the latter containing many undead neurons (considered in more detail below). The remaining cells in our datasets mostly represent sensillar support cells (Figure 1C and Figure S1D), suggesting that *peb-Gal4* labels both neuronal and non-neuronal branches of the SOP lineages; the latter cell types can be investigated in future studies.

We first annotated neurons within these datasets by developmental phase (Figure 1D and Figure S2). All “branches” of cell clusters comprised a continuum through early-, mid- and late-pupal stages (Figure 1D), each presumably reflecting the development of different neuronal lineages. Each phase has distinct transcriptional profiles (Figure S2): early-pupal stage neuronal markers were enriched for genes involved in translation (likely reflecting enhanced protein synthesis capacity); mid-pupal stage neuronal markers were enriched in genes involved in signaling, cell-adhesion, axonogenesis and ion transport (concordant with the wiring of antennal neurons in the brain (Jefferis and Hummel, 2006)); late-pupal stage neurons expressed higher levels of genes involved in ion transport and synaptic transmission (consistent with mature cell functions in neuronal signaling). Cells from mid-developmental stages appeared to be the most transcriptionally divergent between lineages, with late-stage neurons converging to a more similar gene expression profile (as noted previously (McLaughlin et al., 2021)) (Figure 1D).

### High-resolution annotation of the developing antennal sensory neurons

We subclustered these lineages at high resolution, annotating many clusters based on the (chemo)sensory receptor gene(s) expressed in cells of the control dataset (Figure S3). To annotate cells from earlier developmental stages – which mostly lack expression of a diagnostic receptor gene – we used an iterative, retrograde annotation method (see Methods). In brief, marker genes were extracted for each individual neuron cluster – and/or groups of neurons housed in the same sensillum (“sensillar markers”) – and used to identify and annotate additional clusters. These clusters were used as sources of additional earlier marker genes (see example iterations in Figure S4). Ultimately, we could annotate ∼90% of neurons (Figure 1E); most of the remaining cells correspond to the earliest time points, which were difficult to distinguish transcriptionally. A subset of these early cells expresses glial markers (Figure 1E and Figure S3B); these might correspond to the Nbb-derived glia within the neuronal lineages (Endo et al., 2007; Rodrigues and Hummel, 2008; Sen et al., 2005). The ∼19,000 annotated neurons in our control atlas represent more than 15-fold coverage of this sensory organ (∼1200 neurons/antenna) (Grabe et al., 2016; Schlegel et al., 2021).

Our annotation of sensory neurons allowed us to document the complements of cell adhesion molecules, neurotransmitter receptors and ion channels for individual neuron types (Figure S5), substantially extending previous analyses (McLaughlin et al., 2021). Such information might point to additional molecules defining the specific anatomical and functional properties of different sensory channels. In the context of understanding the development of the olfactory system, we were also able to extract markers for all sensillar classes (Figure 1F and Figure S6), and those distinguishing the four neuron precursor types across essentially all lineages (Figure 1G and Figure S7). The latter were enriched for genes encoding neural guidance molecules (Figure S7C), concordant with the segregation of the OSNs from different precursor types in the antennal lobe (Endo et al., 2007).

Our atlas encompasses all documented antennal sensory channels (Benton et al., 2025). This contrasts with previous single cell/nuclear RNA-sequencing (sc/snRNA-seq) analyses of the *D. melanogaster* developing antenna, which were able to match only about one-third of known OSN types across three time points (24 h APF, 48 h APF and adult) (Li et al., 2020; McLaughlin et al., 2021) due to limited cell numbers. To further assess the completeness of our dataset, we compared the relative abundance of each sensory neuron class in the control dataset with those expected by *in situ* analysis (typically using RNA FISH or sensory receptor promoter reporter lines) (Benton et al., 2025). These values exhibited a remarkably strong linear relationship, indicating the highly quantitative nature of cell representation in our datasets (Figure 1H).

### Identifying transcription factors essential for lineage-specific development

As a first assessment of the predictive ability of our developmental atlas and to understand how precision in the olfactory map arises, we examined transcription factors (TFs). Across pupal stages each neuronal population expresses a unique, though often highly overlapping, combination of 138 TF genes (Figure S8). We tested the functional relevance of TF expression through study of three very narrowly-expressed TF genes, which we reasoned might have selective and non-redundant roles: *lozenge* (*lz*), encoding an AML/Runt family TF, which is expressed exclusively in ab10 Or67a and Or85f neurons, and the tandemly-organized paralogs *ladybird early* (*lbe*) and *ladybird late* (*lbl*), encoding NK-like homeobox TFs, which are co-expressed in Or23a and Or83c neurons in ai2 (Figure 2A and Figure S8).

**Figure 2.**
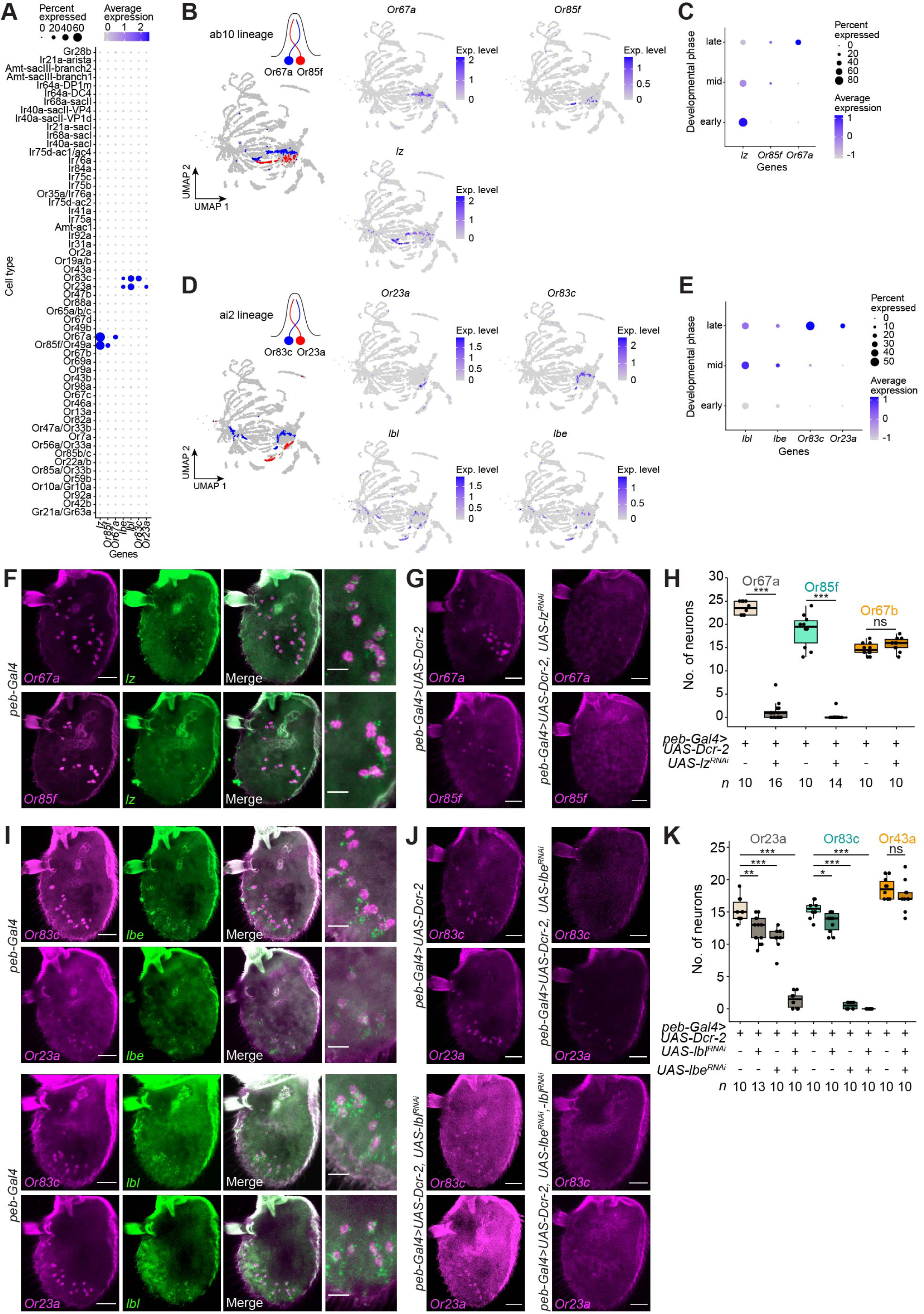
Novel lineage-specific transcription factors. **(A)** Expression of the TFs *lozenge* (*lz*), *ladybird early* (*lbe*) and *ladybird late* (*lbl*) as well as *Or85f*, *Or67a*, *Or83c* and *Or23a* across cell types (control dataset, except dying lineages, here and in other panels). Expression levels (here and elsewhere) have arbitrary units (see Methods). **(B)** UMAPs highlighting the ab10 lineages (schematized in the cartoon) (left) and the expression of the indicated genes (right). **(C)** Expression of the indicated genes in ab10 cells grouped by developmental phases. **(D)** UMAPs highlighting the ai2 lineages (schematized in the cartoon) (left) and the expression of the indicated genes (right). **(E)** Expression of the indicated genes in ai2 cells grouped by developmental phases. **(F)** RNA FISH on whole-mount antennae of control (*peb-Gal4*) animals with the indicated probes (*n* = 10-16 antennae). Scale bars, 25 µm (or 10 µm for single confocal Z-slice, high-magnification images on the right), here and in other panels. **(G)** RNA FISH on whole-mount antennae of control (*peb-Gal4,UAS-Dcr2*) and *lz^RNAi^* (*peb-Gal4,UAS-Dcr-2/+;UAS-lz^RNAi^/+;*) animals with probes targeting the indicated transcripts. **(H)** Quantification of experiments in (G), together with a control *Or67b* probe (images not shown). Here and elsewhere, box plots illustrate individual data points overlaid on boxes showing the median (thick line), first and third quartiles, while whiskers indicate data distribution limits. *n* is indicated underneath. *** = P < 0.001; ns = P > 0.05, respectively, *t* test. **(I)** RNA FISH on whole-mount antennae of control (*peb-Gal4*) animals with the indicated probes (*n* = 10-16 antennae). **(J)** RNA FISH experiment on whole-mount antennae of control (*peb-Gal4,UAS-Dcr-2*), *lbl^RNAi^* (*peb-Gal4,UAS-Dcr-2/+;;UAS-lbl^RNAi^/+*), *lbe^RNAi^* (*peb-Gal4,UAS-Dcr-2/+;UAS-lbe^RNAi^/+;*), and *lbl^RNAi^,lbe^RNAi^* (*peb-Gal4,UAS-Dcr2/+;UAS-lbe^RNAi^/+;UAS-lbl^RNAi^/+*) animals with the indicated probes targeting the indicated transcripts. **(K)** Quantification of experiments in (J), together with a control *Or43a* probe (images not shown). *n* is indicated underneath. * = P < 0.05; ** = P < 0.01; *** = P < 0.001; Wilcoxon rank sum test followed by Bonferroni correction for multiple comparisons. ns = P > 0.05, *t* test.

In all cases TF expression precedes that of the *Or* genes (Figure 2B-E). We first validated these predicted expression patterns *in situ*, revealing selective expression of the TFs in the expected neuron populations (Figure 2F and 2I). We knocked down TF expression by using the *peb-Gal4* to drive TF RNAi transgenes (Figure 2G-H and 2J-K). *lz^RNAi^* led to essentially complete loss of expression of *Or67a* and *Or85f*, while not affecting a control population expressing *Or67b*. *lbe^RNAi^* had, respectively, a mild or very strong effect on expression of *Or23a* and *Or83c*. While *lbl^RNAi^* alone barely affected expression of either receptor, it strongly enhanced the *lbe^RNAi^* phenotype, indicating partial redundancy of these TFs in ai2 neurons. This function appears to be specific, as a control receptor Or43a was unaffected (Figure 2K). Together, these data demonstrate an essential and selective role of *lz* and *lbe*/*lbl* in controlling ab10 and ai2 OSN development, respectively.

As *lz* and *lbe/lbl* are common to both OSNs within their respective sensilla, we hypothesize that these TFs act prior to the last lineage division that produces these distinct OSN types, rather than directly in receptor expression. Lbe/Lbl have previously been associated with cell fate specification in muscle and heart cells (Junion et al., 2007), rather than sensory neurons. By contrast, Lz was one of the first TFs implicated in antennal development, with a widespread role in antennal sensillar patterning that presumably occurs at an earlier stage within the antennal imaginal disc (Stocker et al., 1993). Our findings reveal a later, lineage-specific role for Lz, illustrating how TFs can play multiple roles within the development of this sensory system.

### Precision and promiscuity of tuning receptor and co-receptor transcription

The analyses above indicate the comprehensive and functionally-predictive nature of our atlas, setting the stage for exploring how precision in the olfactory system is established. We first examined the spatio-temporal expression patterns of receptor genes (Figure 3A and Figure S9). In early developmental stages, only one gene was detected, the *Ir25a* co-receptor (discussed further below) (Figure S9). By mid-stages, transcripts for over a dozen tuning Irs or Ors in different populations of neurons, as well as *Ir8a* and *Ir93a* co-receptors, were detectable (Figure S9). By late stages, essentially all neuronal populations were reliably expressing a specific tuning receptor gene(s) (Figure 3A and Figure S9). As expected, the majority of these expressed a single tuning receptor, but our data confirmed the known cases of tuning receptor co-expression. The latter include the two subunits of the CO_2_ receptor (*Gr21a/Gr63a*) (Jones et al., 2007; Kwon et al., 2007), cases of co-expression of genomically- and phylogenetically-distant receptor genes (e.g., *Or56a/Or33a*), and several examples of co-expression of closely-related, tandemly-arranged, receptor paralogs (e.g., *Or19a/Or19b*, *Or22a/Or22b*), where genes in these clusters likely retain conserved *cis*-regulatory sequences after gene duplication. We note that co-occurrence of *Ir75a*, *Ir75b* and *Ir75c* transcripts does not reflect co-expression of these genes, but rather runaway transcription in this cluster; protein-coding transcripts are expressed in distinct populations of OSNs (Mika et al., 2021; Prieto-Godino et al., 2017). While both Gr21a and Gr63a are essential for CO_2_ responses (Jones et al., 2007; Kumar et al., 2020; Kwon et al., 2007), it is unclear whether the other cases of co-expression are functionally significant. Co-expressed paralogs might be functionally redundant or represent a transient evolutionary state as one paralog undergoes pseudogenization or neofunctionalization (Auer et al., 2022).

**Figure 3.**
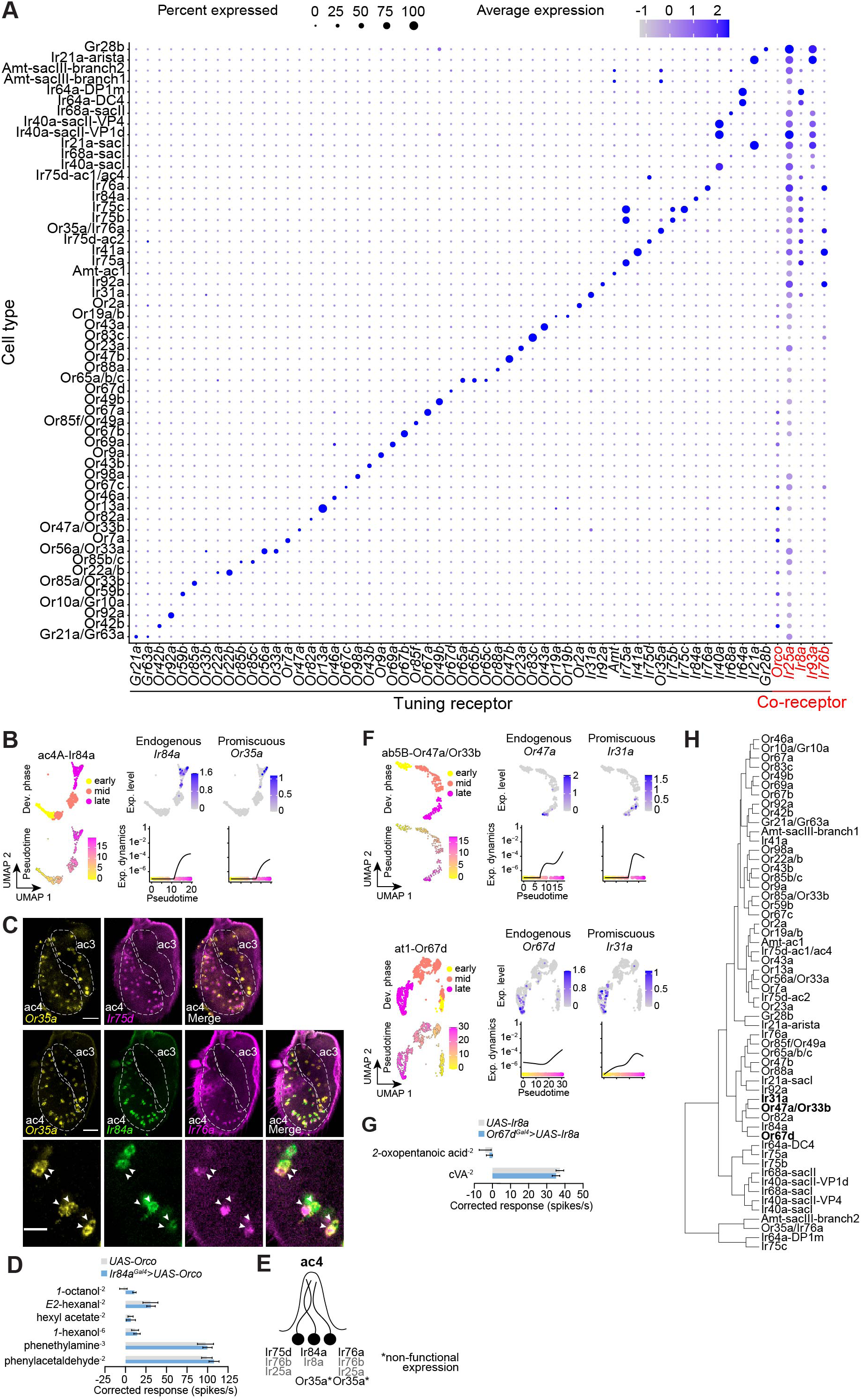
Precision and promiscuity of sensory receptor transcription. **(A)** Dot plot illustrating the expression of tuning receptor and co-receptor subunits across sensory neuron types in the late-stage, control dataset (*peb-Gal4/+;;UAS-unc84:GFP/+*). Only three expected tuning receptors were not detected: (i) *Or49a* (which was, however, detectable in the expected cells – i.e., co-expressed with *Or85f* – in the non-normalized data (not shown), (ii) co-expressed *Or10a* and *Gr10a* (likely because the genes have identical annotations, resulting in reads being filtered out due to unintended mapping to two distinct genes), and (iii) Or33b, which were not detected in Or85a neurons as reported (Fishilevich and Vosshall, 2005), but instead detected in a very small fraction of Or56a/Or33a neurons, potentially reflecting strain specificity. **(B)** Top: UMAPs of the ac4A lineage illustrating the developmental stages and receptor expression patterns (control dataset). Bottom: a pseudotime UMAP and corresponding receptor expression dynamics. **(C)** RNA FISH on whole-mount antennae of control (*peb-Gal4*) animals with probes targeting the indicated transcripts (*n* = 10-12). The ac3 and ac4 sensilla zones are indicated in the top and middle rows. Bottom: ac4A neurons co-expressing *Ir84a* and *Or35a* and ac4B neurons co-expressing *Ir76a* and *Or35a* in a single confocal Z-slice. Scale bars, 25 µm (top and middle rows) or 10 µm (bottom row). **(D)** Electrophysiological responses to the indicated ligands in ac4 sensilla from antennae of control animals (*UAS-Orco*) and animals overexpressing *Orco* in ac4A neurons (*Ir84a^Gal4^/UAS-Orco*). Solvent-corrected responses (mean ± SEM) are shown; see Data S1 for raw data and statistical analyses. **(E)** Schematic summarizing tuning receptor (black) and co-receptor (grey) subunits expressed in ac4 neurons. **(F)** UMAPs of the ab5B (top rows) or at1 (bottom rows) lineages illustrating the developmental stages and receptor expression patterns (control dataset), pseudotime UMAPs and corresponding receptor expression dynamics. **(G)** Electrophysiological responses to the indicated ligands in at1 sensilla from antennae of control animals (*UAS-Ir8a*) and animals overexpressing *Ir8a* in at1 neurons (*Or67d-Gal4/UAS-Ir8a*). Solvent-corrected responses (mean ± SEM) are shown; see Data S1 for raw data and statistical analyses. **(H)** Hierarchical clustering of antennal neuron types based upon differentially expressed TFs (from Figure S8).

Our analyses revealed several cases of unexpected receptor expression. For example, *Or35a* is weakly expressed in ac4 Ir84a OSNs (Figure 3B), in addition to its well-described expression in ac3 neurons (Figure 3A). We confirmed these transcriptomic data *in situ*, detecting *Or35a* in ac4 sensilla, both in a subset of Ir84a neurons, as well as some Ir76a neurons (suggesting it was turned on in these cells at a later time point than we have profiled transcriptionally) (Figure 3C). Here, Or35a is unlikely to be functional, as Orco protein is not expressed in these neurons (Figure 3A); indeed, these neurons do not respond to ligands that activate Or35a neurons in ac3 (Silbering et al., 2011; Yao et al., 2005). We tested whether Or35a has the potential to be functional in Ir84a OSNs through transgenic expression of Orco in these cells, but this did not produce responses to Or35a-dependent ligands (Figure 3D), suggesting that *Or35a* expression is too low, or that the transcript is aberrantly spliced (Shang et al., 2024), and/or that other factors are required for Or35a function (Figure 3E). *Or35a* transcripts were also detected in neurons of sacculus chamber III (sacIII) (Figure 3A and Figure S10A-B), which correspond to the Amt-expressing ammonia-sensing neurons (Vulpe et al., 2021); here it is also unlikely to be functional as these cells do not express Orco (Figure 3A).

Another example of more promiscuous expression was observed for *Ir31a*, which was detected in both Or47a and Or67d neurons, in addition to its own population (Figure 3F). This expression is not detected *in situ* (Benton et al., 2009; Silbering et al., 2011) and is unlikely to be functional as the essential Ir8a co-receptor is not expressed in these neurons (Figure 3A), nor have they been described to respond to Ir31a-dependent ligands (de Bruyne et al., 2001; Munch and Galizia, 2016; van der Goes van Naters and Carlson, 2007). Mis-expression of Ir8a in Or67d neurons also failed to confirm sensitivity to the best known Ir31a agonist, *2*-oxopentanoic acid (Figure 3G). Closer examination of the transcriptomic data revealed that *Ir31a* expression is transient, reaching highest levels prior to the branch termini in these lineages; this pattern contrasts with *Or47a* and *Or67d* expression, which peak at the end of the branches. We wondered whether this transient “ectopic” expression of an *Ir* in these Or neurons reflected similarity in the gene regulatory networks of these distinct cell types. Using information of differentially-expressed TFs (Figure S8), we performed hierarchical clustering analysis of all OSN classes (Figure 3H). Notably, Ir31a, Or47a and Or67d neurons tend to cluster within this tree, raising the possibility that *Ir31a* “eavesdrops” on the complement of TFs of Or47a and Or67d neurons to become transiently activated during development.

We extended our survey to receptors expressed in other chemosensory organs (Figure S11). Amongst Ors, we detected transcripts of the maxillary palp receptor *Or42a* in Ir31a neurons (Figure S11 and S10C). Again, this is unlikely to be functional as Orco is not expressed in these cells (Figure 3A) (Benton et al., 2009), and these neurons do not respond to Or42a ligands (Silbering et al., 2011). Several *bona fide Gr*s were detected in a number of antennal cell types, consistent with observations from previous transcriptomic and transgenic studies (Figure S11 and S10D-E) (Fujii et al., 2015; McLaughlin et al., 2021; Menuz et al., 2014). The functional significance, if any, of *Gr*s in the antenna in unclear (Pal Mahadevan et al., 2022); we favor a hypothesis that such *Gr* expression merely reflects promiscuous transcription of these genes, possibly due to overlap of the set of TFs present in these antennal neurons and the gustatory neurons in which these receptors are normally expressed.

Finally, our datasets confirm the previous observations of broad and partially overlapping expression of various co-receptor genes (Figure 3A) (Abuin et al., 2011; McLaughlin et al., 2021; Task et al., 2022). Whether co-receptors function in every neuron in which they are expressed is unclear. Genetic and electrophysiological analyses indicate an olfactory requirement for Ir8a only with selectively-expressed acid-sensing tuning Irs (e.g., Ir31a, Ir64a), and Ir25a/Ir76b together only with amine-sensing Irs (e.g., Ir41a, Ir76a) (Abuin et al., 2011; Vulpe and Menuz, 2021), despite broader expression all three of these co-receptor genes (Figure 3A) (Abuin et al., 2011; McLaughlin et al., 2021; Task et al., 2022). Loss of Ir25a has been described to affect Or neuron sensitivity to stimuli, with mild increase or decrease of responses, depending upon the neuron and the odor (Task et al., 2022), but the mechanistic basis for such a variable effect on Or neuron responses is unclear. To our knowledge there is only one case of a neuron (in ac3) that contains a functional complement of tuning and co-receptor Ors and Irs (Benton et al., 2025). We suggest that contributions of broadly-expressed co-receptors is constrained by the selective expression of partner tuning subunits: just as tuning receptors without co-receptors are likely to be non-functional, co-receptors without partners might have no or minimal sensory contributions.

### Heterogeneous life and death fates of specific OSN populations

We next turned our attention to the neurons that are normally removed by PCD during antennal development. While such cells can develop into functional sensory neurons when PCD is blocked (Prieto-Godino et al., 2020), we know little about their molecular and developmental properties. To identify cells in the atlas corresponding to those that undergo PCD, we examined the expression of the pro-apoptotic genes *reaper* (*rpr*), *grim*, *sickle* (*skl*) and *head involution defective* (*hid*) as one or more of these genes are transcriptionally upregulated prior to PCD in many developmental contexts (Pinto-Teixeira et al., 2016) (Figure 4A). Indeed, we detected higher expression of *rpr*, *grim* and *skl* in the PCD-blocked dataset than the control dataset, with largely overlapping expression patterns of these genes in a subset of cell clusters representing putative undead neurons (Figure S12A-B). By contrast, *hid* was expressed at similar levels in both PCD-blocked and control datasets (Figure S12A-B), indicating that this gene is not a selective marker of cells that are fated to die (as noted in other tissues (Pinto-Teixeira et al., 2016)), and was therefore discounted as a marker. We therefore combined expression enrichment of *rpr*, *grim* and *skl* into a single “RGS score”, as a quantitative measure of the likelihood that a cell type was fated to die (Figure 4B-C).

**Figure 4.**
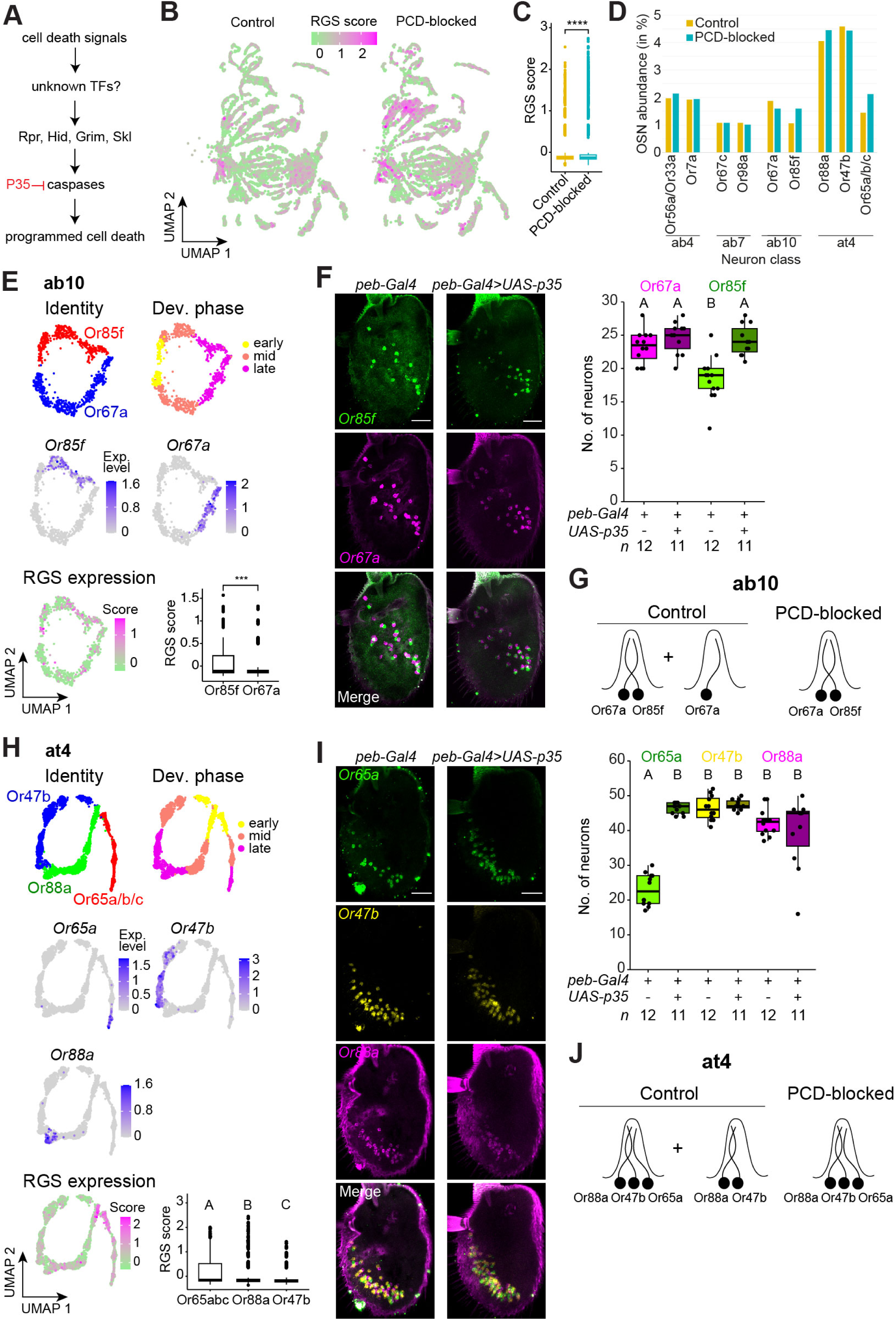
Heterogeneous life and death fates of specific OSN populations. **(A)** Schematic of the programmed cell death cascade in *D. melanogaster*. **(B)** UMAPs illustrating the combined expression levels of the pro-apoptotic genes *reaper* (*rpr*), *grim* and *sickle* (*skl*) – quantified as a “RGS” score – in control (*peb-Gal4/+;;UAS-unc84:GFP/+*) and PCD-blocked (*peb-Gal4/+;UAS-p35/+;UAS-unc84:GFP/+*) datasets. **(C)** RGS score in control and PCD-blocked datasets. Here and elsewhere, boxes show the median (thick line), first and third quartiles, while whiskers indicate data distribution limits. **** = P < 0.0001, Wilcoxon rank sum test. **(D)** Abundance of ab4, ab7, ab10 and at4 neuronal classes in control (yellow) and PCD-blocked (blue) datasets, calculated as the percentage of nuclei for each class relative to the total number of nuclei (from 36 h APF, excluding those forming new clusters in the PCD-blocked dataset). **(E)** Reconstruction of the ab10 sensillum development in the integrated control and PCD-blocked datasets. Cells were extracted from the datasets using sensillar markers (Figure 1F and Figure S6), and reclustered (see Methods); note in these UMAPs we mask a cell cluster that is unique to the PCD-blocked dataset, which is considered further in Figure S14A-C. UMAPs indicate the identity and the developmental phases (top), the expression of olfactory receptors (middle) and RGS score within the lineage (bottom). Boxplot indicates the ranked (left > right) RGS score in ab10 neurons (**** = P < 0.0001, Wilcoxon rank sum test). **(F)** RNA FISH on whole-mount antennae of control (*peb-Gal4*) and PCD-blocked animals (*peb-Gal4/+;UAS-p35/+*) with probes targeting the indicated transcripts. Scale bars, 25 µm. Quantifications are shown on the right. **(G)** Schematic of the inferred subtypes of ab10 sensilla in control and PCD-blocked datasets. **(H)** Reconstruction of the at4 sensillum development in the integrated control and PCD-blocked datasets. UMAPs indicate the identity and the developmental phases (top), the expression of olfactory receptors (middle) and RGS score within the lineage (bottom). Boxplot indicates the ranked (left > right) RGS score in at4 neurons. **(I)** RNA FISH on whole-mount antennae of control (*peb-Gal4*) and PCD-blocked animals (*peb-Gal4/+;UAS-p35/+*) with probes targeting the indicated transcripts. Scale bars, 25 µm. Quantifications are shown on the right. **(J)** Schematic of the inferred subtypes of at4 sensilla in control and PCD-blocked datasets. **(F, H and I)** A, B and C indicate significant differences: P < 0.05 in pairwise comparisons (Wilcoxon rank sum test followed by Bonferroni correction for multiple comparisons).

We next decomposed the atlas of integrated control and PCD-blocked datasets (Figure 1E) into individual sensilla, using sensillar markers (Figure 1F and Figure S6) and compared cells from each dataset. We reasoned that some undead neurons might form cell clusters unique to the PCD-blocked dataset, as investigated in the next section. However, as many undead neurons express receptors characteristic of normal populations of OSNs (Prieto-Godino et al., 2020), we first asked whether any such cells are embedded in clusters of these normal cells, thereby simply creating a larger cluster in the PCD-blocked dataset. Given the highly quantitative representation of cell populations in our antennal atlas (Figure 1H), to identify such undead neurons, we first compared the relative proportion of each neuronal type in control and PCD-blocked datasets (Figure S12C and Figure 4D). While neurons housed in the same sensillum generally displayed similar representations within and across datasets (e.g., Or56a/Or33a and Or7a neurons in ab4, or Or67c and Or98a neurons in ab7, Figure 4D), in several cases in the control dataset we observed different proportions of co-housed neurons. For example, in ab10 Or85f neurons are underrepresented compared to Or67a neurons, and in the trichoid sensillum at4 Or65a/b/c neurons are less abundant than Or47b and Or88a neurons (Figure 4D). Importantly, these mis-matched neuronal representations were at least partially re-equilibrated in the PCD-blocked atlas (Figure 4D and Figure S12C), suggesting individual cell types within a subset of sensilla are developmentally lost due to death.

Focusing first on ab10, we investigated this possibility initially by comparing the pro-apoptotic gene RGS score within individual neuron lineages (Figure 4E). We observed that the Or85f branch has a higher score, consistent with the occurrence of PCD in a subset of these cells. In agreement with these transcriptomic data, RNA FISH for these receptor transcripts revealed a lower number of Or85f neurons compared to Or67a neurons in the antenna; while all of the former are paired with the latter, we detected several cases of isolated Or67a neurons in control animals, but not when PCD was blocked (Figure 4F-G).

Next, we examined at4 (Figure 4H). Here, the underrepresented Or65a/b/c neurons display a higher RGS score during their lineage development compared to Or47b and Or88a lineages. *In situ*, we detected many fewer Or65a neurons than Or47b and Or88a; while the latter two neuron types were always paired, only a subset formed a triplet with Or65a neurons in control animals (Figure 4I-J). These observations match those from electron microscopic studies describing the existence of at4 sensilla housing only two neurons (Nava Gonzales et al., 2021). In PCD-blocked antennae, only the number of Or65a neurons was increased – and all were closely associated with Or47b and Or88a neurons – indicating a subset of these neurons are also naturally removed by PCD (Figure 4I-J). While these *in situ* data indicate 1:1:1 correspondence, we note that within the PCD-blocked dataset, Or65a/b/c neurons were still underrepresented (Figure 4D), suggesting that this dataset lacks annotation of some undead neurons (discussed further below).

Together these observations indicate that PCD acts within several sensilla lineages to selectively remove a subfraction of OSNs normally considered to represent a fully surviving lineage (as opposed to the lineages that are fated to die, which we consider in the next section). It is surprising that such heterogeneity has not been reported previously. In part this might reflect the relatively limited data co-visualizing neurons within the same sensilla in whole-mount antennae – as opposed to cryosections (e.g., (Couto et al., 2005)) – which is essential to view the pairing patterns of the entire cell population. Additionally, it is possible that during electrophysiological recordings, sensilla that do not have the “expected” numbers of neurons are disregarded from detailed study. The functional significance of sensillum heterogeneity is unclear. While the lack of specific neurons in sensilla would eliminate the ephaptic inhibition that can occur between co-housed neurons (Su et al., 2012; Zhang et al., 2019), the loss of these cells might not necessarily be adaptive, as discussed below.

### Diverse states of neurons fated to die during antennal development

We also discovered distinct types of undead neurons represented by cell branches in the developmental trajectory of several sensilla that were present only in the PCD-blocked dataset. For example, in ac3I/II sensilla – housing Or35a and either Ir75b (ac3I) or Ir75c (ac3II) neurons (the lineages of these two subtypes could not be fully distinguished and were considered together) – we observed two extra branches that were extinguished around 30-36 h APF in control antennae but maintained in late pupae in PCD-blocked antennae (Figure 5A). These clusters have a high RGS score, supporting their classification as undead neurons (Figure 5B). One cluster displays a signature of Naa precursor type and the other tentatively Nbb, complementing the assigned Nab and Nba identities of Ir75b/Ir75c and Or35a neurons, respectively (Figure 5C). We surveyed chemosensory receptors in these clusters, finding that the undead Naa neuron expresses *Ir75d* (Figure 5D) – as well as *Ir* co-receptors (Figure S13A) – similar to the expression of *Ir75d* in Naa neurons in ac1, ac2 and ac4 sensilla in wild-type antennae (Benton et al., 2025; Endo et al., 2011). We validated these transcriptomic data *in situ* by demonstrating the pairing of an *Ir75d*-expressing neuron with Ir75b/Ir75c neurons when PCD was blocked but not in control antennae (Figure 5E-G). By contrast, the undead, putative Nbb neuron did not detectably express any tuning receptor, although we did observe the expression of multiple *Ir* co-receptors in these cells (Figure S13A).

**Figure 5.**
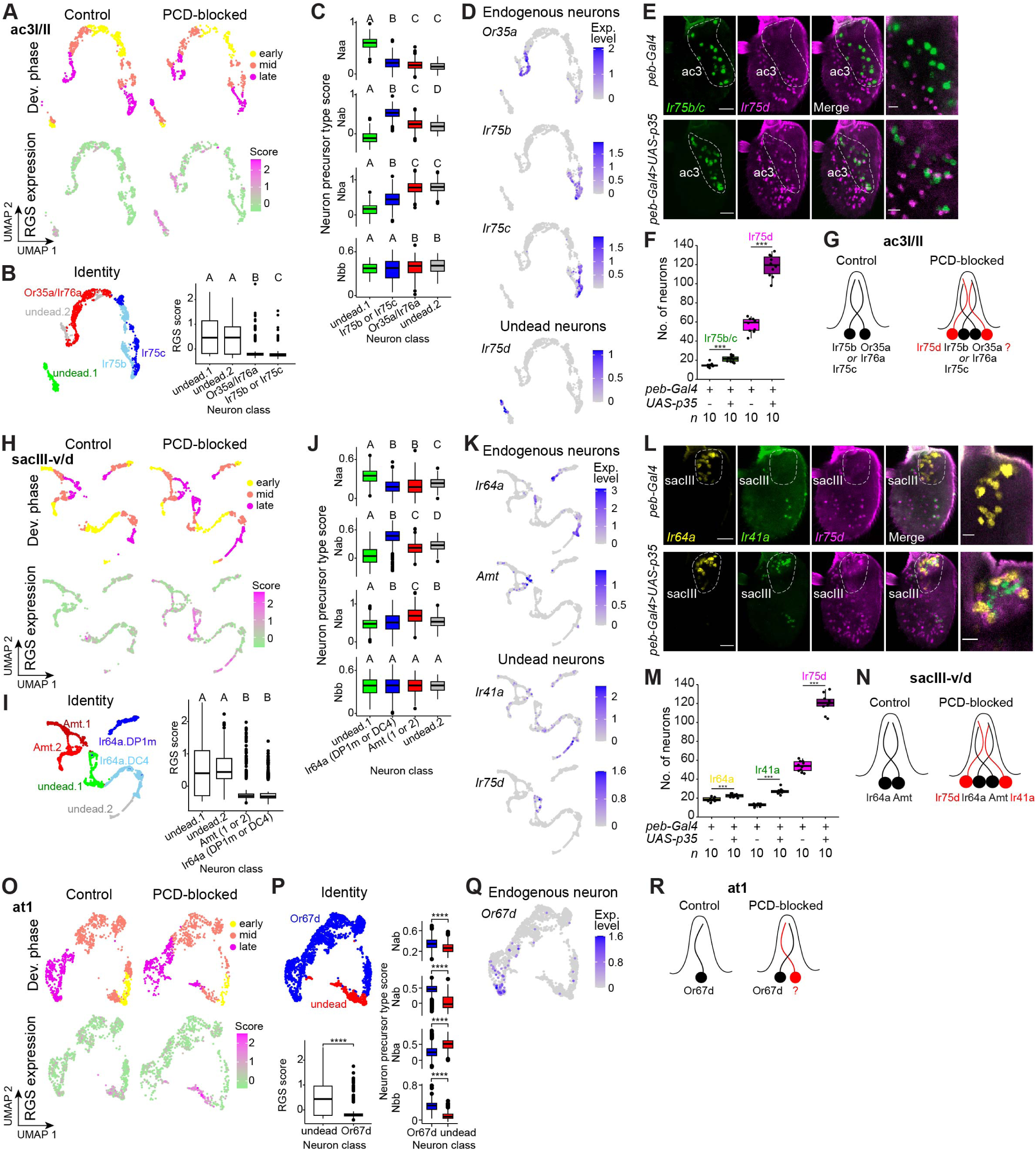
Identification of novel populations of undead neurons. **(A)** UMAPs of the ac3I/II lineages in control and PCD-blocked datasets illustrating the developmental stages (top) and RGS expression (bottom), revealing cells with high RGS score that are present exclusively in PCD-blocked animals. **(B)** Left: lineage annotation of ac3I/II sensilla in integrated control and PCD-blocked data. Right: ranked RGS scores. **(C)** Neuron precursor type score for each OSN in the ac3I/II integrated dataset. **(D)** UMAPs of the integrated ac3I/II dataset illustrating the expression of sensory receptors in endogenous and undead neurons. **(E)** RNA FISH on whole-mount antennae of control (*peb-Gal4*, top row) and PCD-blocked animals (*peb-Gal4/+;UAS-p35/+;* bottom row) with probes targeting the indicated transcripts. The ac3 zone is indicated. Right column images show a higher magnification of single confocal Z-slice in the ac3 zone. Scale bars, 25 µm (left images) or 10 µm (right images). **(F)** Quantification of OSN numbers from the experiments in (E) (*n* is indicated underneath). *** indicates P < 0.001, *t* test. **(G)** Schematic of the inferred types of ac3I/II sensilla in control and PCD-blocked antennae. **(H)** UMAPs of the sacIII-v/d lineages in control and PCD-blocked datasets illustrating the developmental stages (top) and RGS expression (bottom), revealing cells with high RGS score that are present exclusively in PCD-blocked animals. **(I)** Left: lineage annotation of sacIII-v/d sensilla in integrated control and PCD-blocked data. Right: ranked RGS scores. **(J)** Neuron precursor type score for each OSN in the sacIII-v/d integrated dataset. **(K)** UMAPs of the integrated sacIII-v/d dataset illustrating the expression of sensory receptors in endogenous and undead neurons. **(L)** RNA FISH on whole-mount antennae of control (*peb-Gal4*, top row) and PCD-blocked animals (*peb-Gal4/+;UAS-p35/+;* bottom row) with probes targeting the indicated transcripts. The sacIII zone is indicated. Right column images show a higher magnification of single confocal Z-slice in the sacIII zone. Scale bars, 25 µm (left images) or 10 µm (right images). **(M)** Quantification of OSN numbers from the experiments in (L) (*n* is indicated underneath). *** indicates P < 0.001, *t* test. **(N)** Schematic of the inferred states of sacIII-v/d sensilla in control and PCD-blocked antennae. **(O)** UMAPs of the at1 lineages in control and PCD-blocked datasets illustrating the developmental stages (top) and RGS expression (bottom), revealing cells with high RGS score that are present exclusively in PCD-blocked animals. **(P)** Top: lineage annotation of the at1 sensillum in control and PCD-blocked integrated data. Bottom: ranked RGS scores (left) and precursor type scores (right). **** indicates P < 0.0001, Wilcoxon rank sum test. **(Q)** Endogenous expression of *Or67d* in the control and PCD-blocked integrated data. No receptor was robustly detected in the undead neuron population. **(R)** Schematic of the inferred states of the at1 sensillum in control and PCD-blocked antennae. **(B,C,I,J)** A-D letters indicate significant differences: P < 0.05 in pairwise comparisons (Wilcoxon rank sum test followed by Bonferroni correction for multiple comparisons).

Another case of undead neuron clusters was found in sacIII sensilla, which normally house Ir64a and Amt neurons. We again observed two additional cell clusters in the PCD-blocked dataset with elevated RGS scores (Figure 5H-I). Here, one Naa cluster expressed *Ir75d*, while in the other (of unclear precursor type) we detected *Ir41a* (Figure 5J-K). In both cases, we also detected the corresponding *Ir* co-receptors (Figure S13B). We confirmed *in situ* the presence of Ir75d and Ir41a neurons neighboring Ir64a neurons in sacIII in PCD-blocked antennae, but not in wild-type antennae (Figure 5L-N). The transcriptomic data suggested that a subset of the undead Ir75d neurons might also express *Ir64a* (Figure 5K), but we did not observe clear co-expression *in situ* (Figure 5L).

In several additional sensilla classes, we observed undead neuron populations that did not detectably express any receptors (at least up to 80 h APF). In at1, in addition to the *Or67d*-expressing neuron, we detected a second, Nba-derived neuron in the PCD-blocked dataset (Figure 5O-R) consistent with the previous electrophysiological detection of a second neuron (Prieto-Godino et al., 2020). In ab10 in PCD-blocked antennae – beyond the increase in Or85f neuron numbers described above – we detected an extra neuron of unclear precursor type (Figure S14A-D). Finally, in ab5 and sacI, we detected very small populations of likely Naa and Nbb undead neurons, respectively (Figure S14E-L).

Beyond these cases, for the majority of sensillar classes, we did not detect clear evidence for undead neurons (e.g., ai3 and ab4) (Figure S14M-N). This was surprising because, under the canonical model of the SOP lineage (Chai et al., 2019; Endo et al., 2007; Endo et al., 2011), we expected all sensilla to have the potential to produce four terminal cells. This might reflect a technical artefact, for example, a failure to efficiently block PCD in all lineages. It is also possible that such undead cells were simply not recognized as belonging to specific sensilla. This is the case for any Nbb-derived glia (Endo et al., 2007; Rodrigues and Hummel, 2008; Sen et al., 2005), which were not considered in our analyses. However, with two exceptions (asterisk in Figure 1E, a second Ir75d neuron in ac4 (described below)), we did not identify additional populations of more mature undead neurons that are not associated with a particular sensillum) or with robust ectopic receptor expression. It is conceivable that the canonical OSN lineage is not universal and that some sensilla housing two neurons result from lack of a final cell division in the lineage, rather than PCD of two of the daughters of such a division.

### Mamo is required to promote programmed cell death of Ir75d neurons

Although PCD is the most common fate of sensory neuron precursors – if we consider death as a single fate across all sensillum classes – we know essentially nothing about how it is stereotypically specified. Our data indicate that transcriptional activation of *rpr*, *grim* and/or *sickle* is likely to be the key inductive step, similar to other tissues (Sen et al., 2004). However, very little is known about the gene regulatory network upstream of these pro-apoptotic genes in any lineage, and whether these are common to, or distinct between, sensilla. As a first step, we sought TFs required to promote PCD in neurons in specific sensillar types, focusing first on the dying lineages in ac3I/II and sacIII that both express *Ir75d*.

Comparison of the transcriptomes of undead Ir75d neurons in the PCD-blocked dataset with the normal Ir75d neurons housed in ac1, ac2 and ac4 revealed, as expected, *rpr*, *grim* and *skl* to be more highly expressed in the former (Figure 6A and Data S2). The gene displaying the greatest enrichment in the undead Ir75d neurons was *mamo* (*maternal gene required for meiosis*) (Figure 6A)*. Mamo* encodes a zinc finger C2H2 protein, which we hypothesized was a PCD-promoting TF. Consistently, in *mamo^RNAi^* antennae we observed an increase in the number of *Ir75d*-expressing neurons (Figure 6B). These were located both in ac3I/II (paired with *Ir75b/Ir75c* expressing neurons), as well as in sacIII (paired with Ir64a neurons) (Figure 6C-F). These phenotypes were not seen upon RNAi of several other TF genes that have enriched expression in undead Ir75d neurons (Figure 6A and Data S3).

**Figure 6.**
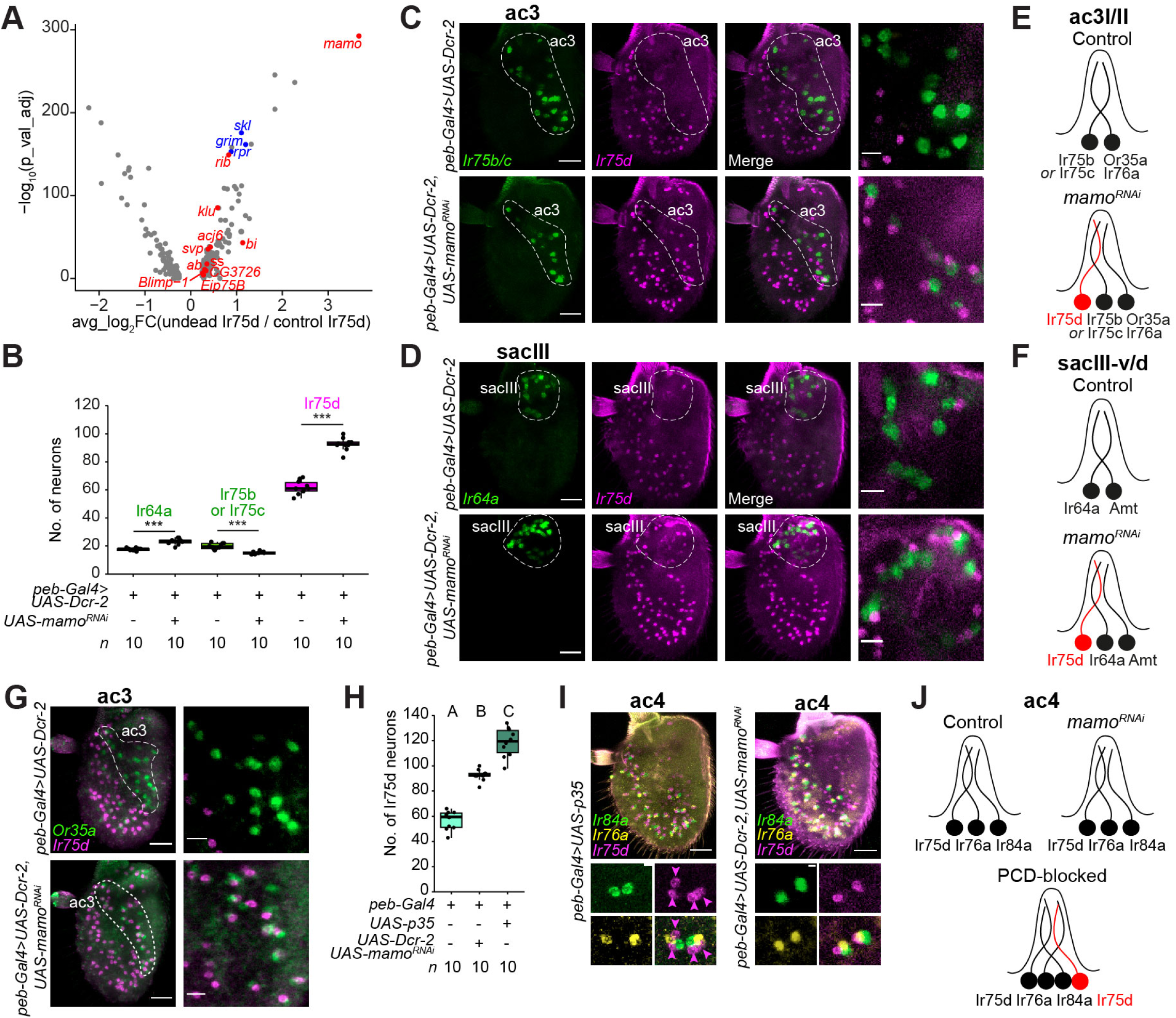
*Mamo* specifies PCD in Ir75d neuron lineages. **(A)** Volcano plot illustrating differentially-expressed genes between undead Ir75d OSNs (in ac3 and sacIII) and normal Ir75d OSNs (in ac1, ac2 and ac4) (pseudobulk analysis, *x* axis = log_2_FC(“undead”/control), *y* axis = -log_10_(adjusted P), log_2_FC > 0.25, % of positive nuclei > 0.25). Pro-apoptotic genes are highlighted in blue, and TFs whose expression is enriched in the undead Ir75d neurons are highlighted in red. See Data S2 for the top 20 genes enriched in undead and living Ir75d neuron counterparts. **(B)** Quantification of OSN numbers in the indicated genotypes (*n* is indicated underneath). *** indicates P < 0.001, *t* test. **(C,D)** RNA FISH on whole-mount antennae of control (*peb-Gal4,UAS-Dcr-2*, top row) and *mamo^RNAi^* animals (*peb-Gal4,UAS-Dcr-2/+;UAS-mamo^RNAi^/+*, bottom row) with probes targeting the indicated transcripts. ac3 (C) and sacIII (D) zones are shown. Right column images show a higher magnification within the ac3 and sacIII zones of a single confocal Z-slice, showing “undead” Ir75d neurons upon *mamo^RNAi^* in both ac3 and sacIII sensilla. Scale bars, 25 µm (left images) or 10 µm (right images). **(E,F)** Schematic of the inferred states of ac3I/II (E) and sacIII-v/d (F) sensilla in control and *mamo^RNAi^* conditions. **(G)** RNA FISH on whole-mount antennae of control (*peb-Gal4,UAS-Dcr-2;* top) and *mamo^RNAi^* animals (*peb-Gal4,UAS-Dcr-2/+;UAS-mamo^RNAi^/+;* bottom) with probes targeting the indicated transcripts. The ac3 zone is indicated. Right column images show a higher magnification in the ac3 zone in a single confocal Z-slice. Scale bars, 25 µm, or 10 µm for high-magnification images. **(H)** Quantification of Ir75d neurons in the indicated genotypes (*n* is indicated underneath); data is replotted from Figures 5K and 6B, to highlight the mismatch of Ir75d neuron numbers in *mamo^RNAi^* and PCD-blocked animals. A-C letters indicate significant differences: P < 0.05 in pairwise comparisons (Wilcoxon rank sum test followed by Bonferroni correction for multiple comparisons). **(I)** RNA FISH on whole-mount antennae of PCD-blocked (*peb-Gal4/+;UAS-p35/+; left*) and *mamo^RNAi^* animals (*peb-Gal4,UAS-Dcr-2/+;UAS-mamo^RNAi^/+;* right) with probes targeting the indicated transcripts, showing an additional *Ir75d* expressing neuron in ac4. Bottom quadrants show higher magnification of a single Z-slice. Scale bars, 25 µm, or 3 µm for quadrants. **(J)** Schematic of the inferred states of ac4 sensilla in control, *mamo^RNAi^* and PCD-blocked antennae.

The *mamo^RNAi^* phenotype is similar to that of PCD-blocked antennae (Figure 5A-N) but, in principle, loss of this TF could simply lead to ectopic *Ir75d* expression in another neuron type within ac3 and sacIII sensilla. Indeed, Mamo was previously characterized for its role in defining cell fate in the neuroblasts (neural stem cells) of the mushroom body (Liu et al., 2019). However, several observations argue against this possibility in OSNs. First, in *mamo^RNAi^* antennae the Ir75d neuron in ac3 is also paired with the Or35a neuron (Figure 6G), verifying that ac3 sensilla contains distinct neurons expressing Ir75b (or Ir75c), Or35a and Ir75d. Second, loss of Mamo does not appear to affect fate specification of several living OSN populations in which it is expressed (Figure S8), including Or35a, Ir75b, Ir75c and Ir64a neurons, although we noticed modest changes in Ir75b/Ir75c and Ir64a neuron numbers upon *mamo^RNAi^* (Figure 6B). Third, we traced the projections of *Ir75d*-expressing neurons to the antennal lobe using an *Ir75d promoter*-CD4:GFP reporter. In control animals, these neurons converge on the VL1 glomerulus (Figure S15). A similar convergence was observed in both PCD-blocked and *mamo^RNAi^* genotypes (Figure S15), consistent with both types of genetic manipulations producing equivalent undead Ir75d neurons with the same projection properties as normal Ir75d neurons.

Together, these data implicate Mamo as part of the gene-regulatory network inducing cell death of Ir75d neurons in both ac3I/II and sacIII lineages, revealing a novel function of this TF. We note, however, that Mamo is expressed across a large number of OSN populations (including normal Ir75d neurons in ac2) (Figure S8), indicating that this TF is unlikely to be instructive alone for PCD fate but rather functions in a context-dependent manner to promote death. Moreover, there also appear to be Ir75d neurons that die in a *mamo-*independent manner, as inhibition of PCD with P35 led to more Ir75d neurons than in *mamo^RNAi^* antennae (Figure 6H). Reviewing *Ir75d* expression patterns in PCD-blocked antennae (Figure 5E and 5L), we noticed pairs of Ir75d neurons in the ac4 region. Visualizing markers for ac4 (Ir84a and Ir76a), we confirmed the existence of two Ir75d neurons in these sensilla when PCD is blocked but not in *mamo^RNAi^* antennae (Figure 6I-J). Thus, the ac4 lineage has the potential to form a second Ir75d neuron, which is normally fated to die through other, unknown, TFs.

### Slp2 is required to promote programmed cell death of at1 neurons

We next investigated the undead neuron population in at1. As we did not identify a sensory receptor in this neuron to permit comparison with an endogenous cell population, we sought TFs enriched in the undead cell cluster compared with the co-housed Or67d neuron (Figure 7A). The at1 undead neuron exhibited higher expression of the forkhead TF gene *sloppy paired 2* (*slp2*) (Figure 7A-B). To examine the requirement for *slp2* we performed electrophysiological recordings in at1 sensilla in control and *slp2^RNAi^* animals, as we could not visualize the undead neuron through RNA FISH for a receptor transcript. Control at1 sensilla house a single neuron, detected as spikes of a uniform amplitude, corresponding to the cVA-responsive Or67d neuron (Figure 7C-E). By contrast, in a large fraction of *slp2^RNAi^* at1 sensilla, we detected an additional, smaller spike amplitude, indicative of a second, undead neuron, phenocopying the consequences of blocking PCD with P35 (Figure 7C-E).

**Figure 7.**
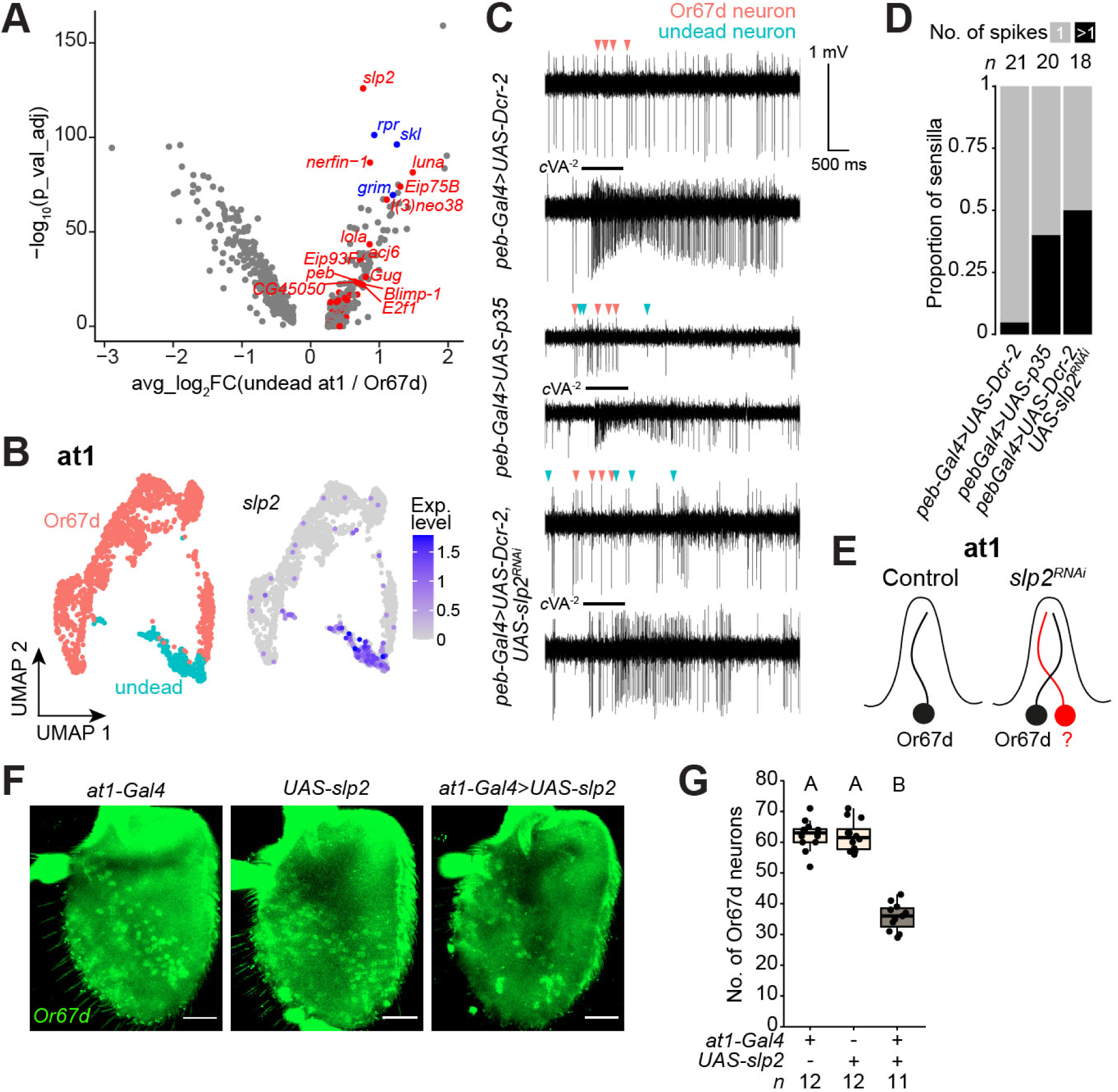
*Slp2* activity is necessary and sufficient to specify PCD in the at1 lineage. **(A)** Volcano plot illustrating differentially-expressed genes between at1 undead and Or67d OSN populations (pseudobulk analysis, *x* axis = log_2_FC(“undead”/control), *y* axis = -log_10_(adjusted P), log_2_FC > 0.25, % of positive nuclei > 0.25). Pro-apoptotic genes are highlighted in blue, and TFs whose expression is enriched in the undead at1 neurons are highlighted in red. **(B)** UMAPs of the at1 lineage (integrated control and PCD-blocked conditions), illustrating neuron identity and *slp2* expression. **(C)** Traces of spontaneous (top) and cVA-evoked (bottom) electrophysiological activity from the at1 sensillum in the antennae of control (*peb-Gal4,UAS-Dcr-2*), PCD-blocked (*peb-Gal4/+;UAS-p35/+;*) and *slp*2*^RNAi^* (*peb-Gal4,UAS-Dcr-2/+;;UAS-slp2^RNAi^/+*) animals. Red and blue arrows indicate spikes of the Or67d and undead neurons, respectively. **(D)** Quantification of the proportion of at1 sensilla housing 1 or >1 spike amplitude in the indicated genotypes (*n* is indicated above). **(E)** Schematic of the inferred states of the at1 sensillum in control and *slp2^RNAi^* antennae. **(F)** RNA FISH experiment on whole-mount antennae of control animals (;;*UAS-slp2-ORF-3HA/+* and ;;*at1-Gal4/+*) and in animals over-expressing *slp2* specifically and transiently in the at1 lineage during development (Chai et al., 2019) (;;*UAS-slp2-ORF-3HA/at1-Gal4*) with probes targeting *Or67d*. Scale bar, 25 µm. **(G)** Quantification of Or67d neurons in the indicated genotypes (*n* is indicated underneath). A and B letters indicate significant differences: P < 0.05 in pairwise comparisons (Wilcoxon rank sum test followed by Bonferroni correction for multiple comparisons).

To further test if *slp2* activity was sufficient to promote PCD, we misexpressed *slp2* in developing Or67d neurons. We used the *at1-Gal4* driver, which is selectively expressed in the at1 lineage from the SOP stage until around 30 h APF, although it only covers about half of the at1 SOPs (Chai et al., 2019). Strikingly, this manipulation led to reduction in Or67d neuron number by around 50% (Figure 7F-G). Together, these data demonstrate that Slp2 activity is necessary and sufficient to promote PCD within the at1 lineage. Similar to Mamo, Slp2 was previously characterized for its role in cell fate diversification within neuroblast divisions, notably as part of the temporal series of TFs controlling optic lobe neuron generation (Konstantinides et al., 2022; Zhu et al., 2022). It thus appears that the antennal SOP lineages co-opt pleiotropic neuronal TFs as part of the gene regulatory networks that promote PCD.

### Context-dependent requirement for Slp2 in promoting programmed cell death

Beyond the undead at1 OSNs, we noticed that *slp2* is expressed in several other populations of OSNs. These include those fated to die (in ab10 and ab5 sensilla), lineages partially eliminated by PCD during antennal development (Or65a/b/c neurons in at4 and Or85f neurons in ab10) as well as in several classes of normal surviving neurons (Or19a/b, Or43a, Or69a, Or88a, Or10a/Gr10a) (Figure 8A). The expression pattern of *slp2* therefore suggested a broader role for this TF in fate specification in the antenna. To test this hypothesis, we surveyed the consequences of *slp2^RNAi^* on other populations of OSNs. Loss of *slp2* had minimal or no effect on Or19a, Or43a and Or69a expression (Figure 8B-C). *slp2^RNAi^* did however lead to increases in ab10 Or85f neurons (Figure 8D-F) and at4 Or65a/b/c (Figure 8G-I) neurons, restoring the 1:1 relationship with other neurons in their respective sensilla; this manipulation phenocopies the effect of PCD inhibition (Figure 4F and 4I). In at4, we note that Or88a neurons express appreciable levels of *slp2* (Figure 8A), but this TF did not appear to have a major role in their fate specification (Figure 8H).

**Figure 8.**
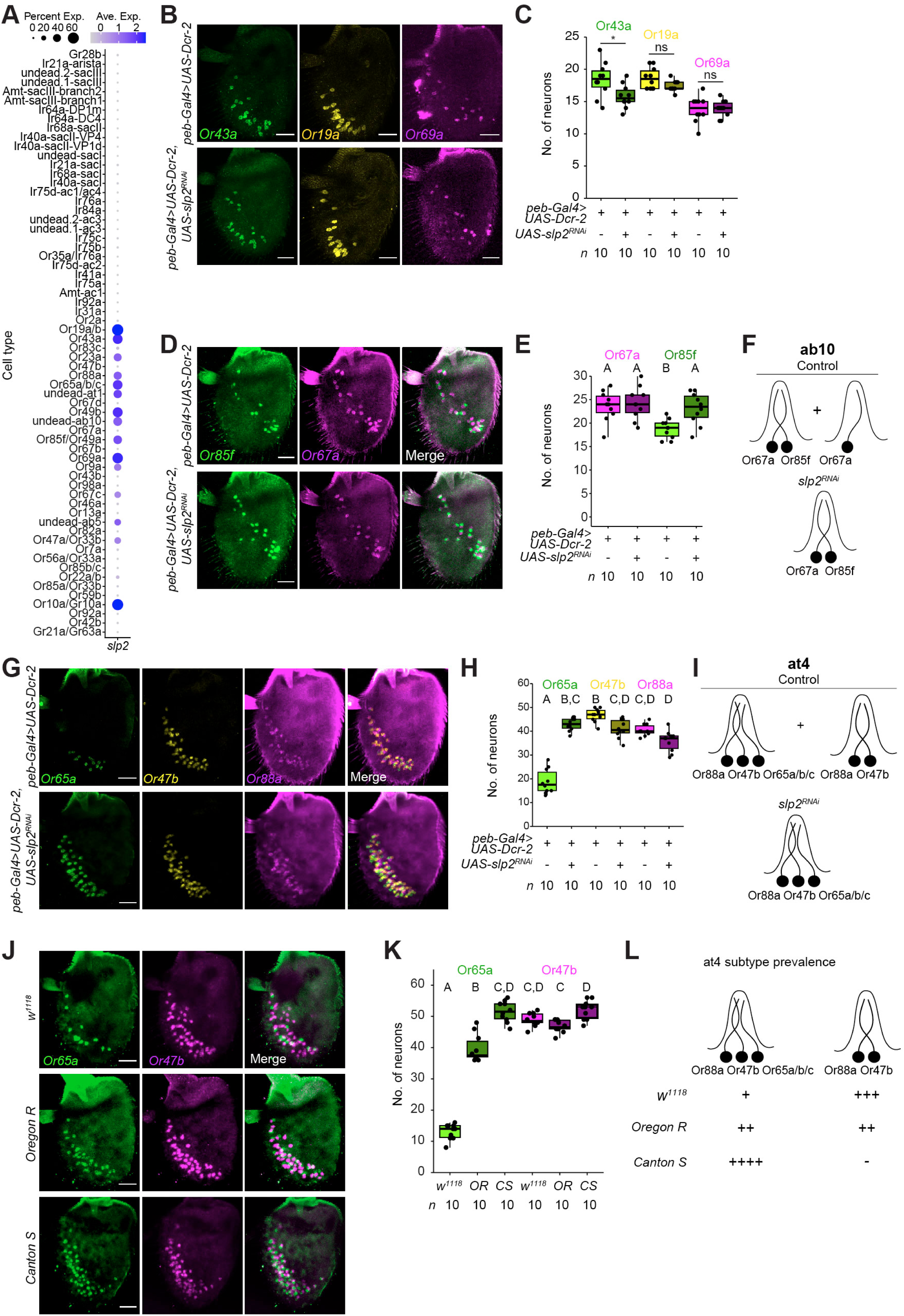
Context-specific requirement for Slp2 in PCD. **(A)** Dot plot showing the expression of *slp2* in all annotated neuron lineages described in this work (control and PCD-blocked integrated datasets). **(B)** RNA FISH on whole-mount antennae of control (*peb-Gal4,UAS-Dcr-2*, top row) and *slp2^RNAi^* animals (*peb-Gal4,UAS-Dcr-2/+;;UAS-slp2^RNAi^/+*, bottom row) with probes targeting the indicated transcripts. Scale bar, 25 µm.xrele **(C)** Quantification of neurons from (B). *n* is indicated below each condition. * and ns indicate P < 0.05 and P > 0.05, respectively, *t* test. **(D)** RNA FISH on whole-mount antennae of control (*peb-Gal4,UAS-Dcr-2*, top row) and *slp2^RNAi^* animals (*peb-Gal4,UAS-Dcr-2/+;;UAS-slp2^RNAi^/+*, bottom row) with probes targeting the indicated transcripts. Scale bar, 25 µm. **(E)** Quantification of neurons from (D). *n* is indicated below each condition. **(F)** Schematic of inferred states of the ab10 sensillum in control and *slp2^RNAi^* antennae. **(G)** RNA FISH on whole-mount antennae of control (*peb-Gal4,UAS-Dcr-2*, top row) and *slp2^RNAi^* animals (*peb-Gal4,UAS-Dcr-2/+;;UAS-slp2^RNAi^/+*, bottom row) with probes targeting the indicated transcripts. Scale bar, 25 µm. **(H)** Quantification of neurons from (G). *n* is indicated below each condition. **(I)** Schematic of inferred states of the at4 sensillum in control and *slp2^RNAi^* antennae. **(J)** RNA FISH on whole-mount antennae of control (*peb-Gal4,UAS-Dcr-2*, top row) and *slp2^RNAi^* animals (*peb-Gal4,UAS-Dcr-2/+;;UAS-slp2^RNAi^/+*, bottom row) with probes targeting the indicated transcripts. Scale bar, 25 µm. **(K)** Quantification of neurons from (G). *n* is indicated below each condition. **(L)** Schematic of inferred states of the at4 sensillum in the antennae of widely used *Drosophila melanogaster* strains. The number of “+” sign reflects the relative prevalence of at4 subtypes (-= absent). **(E,H,K)** A-D letters indicate significant differences: P < 0.05 in pairwise comparisons (Wilcoxon rank sum test followed by Bonferroni correction for multiple comparisons).

These observations argue that the contribution of Slp2 to PCD (or other developmental processes) is context-specific: in some cells (e.g., the dying Nba precursor in at1) it has an essential function, while in others (e.g., Or88a neurons), it has little or no role. at4 Or65a/b/c neurons and ab10 Or85f neurons represent intriguing intermediate cases, as they appear to undergo PCD heterogeneously in a *slp2*-dependent manner. We suggest that such cases of PCD represent “collateral damage” resulting from the expression of Slp2 in these cells that, perhaps due to developmental noise, reaches a minimal threshold of expression to promote PCD in some but not all neurons. In line with this notion, we noticed that different wild-type strains exhibited varying degrees of Or65a/b/c neuron loss: *w^1118^* have (like our *peb-Gal4* strain) low numbers of Or65a/b/c neurons compared to Or47b neurons, Canton-S has an equal number of these two populations, while Oregon R flies have an intermediate number of Or65a/b/c neurons (Figure 8J-L). These observations are consistent with the possibility that this trait is not a fixed, adaptive phenotype of *D. melanogaster*.

## Discussion

The structural and functional properties of neural circuits are often considered as optimized to fulfil their role in controlling animal behavior. In reality, however, these properties represent just a snapshot in evolutionary time, neither precisely the same as in the past, nor necessarily maintained in the future. Understanding the nature of this snapshot in the context of a continuous process of change can offer insights into how nervous systems evolve. The insect olfactory system is a particularly attractive model for studying this phenomenon: this sensory system can be subject to strong environmental selection pressure as the bouquet of external volatile cues changes, and the typically large, rapidly reproducing populations of insect species provide the necessary genetic substrate for evolutionary modifications. Using *D. melanogaster* as a model, we have characterized developmental properties of OSN lineages to reveal features of this species’ olfactory system underlying its functional stereotypy, but also how these offer the potential for evolution.

Through high resolution spatio-temporal transcriptomic profiling of developing neurons, we first confirmed the expected global precision of receptor transcription in the olfactory system, typically a single tuning receptor per neuron type. However, we also reveal that this control is imperfect. We describe several examples of co-expression of tuning receptors of the same or different families. Such co-expression – sometimes transient – presumably reflects similarities in the gene regulatory networks controlling receptor expression in distinct neuronal classes, as suggested by the high degree of overlap in the set of TFs we found in different cell types. The promiscuity in receptor expression is constrained, nevertheless, by the necessity to have the correct complement of tuning receptor and co-receptor subunits to form a functional complex. This seemingly-ectopic expression of receptor genes does permit a degree of evolvability as it is possible that only transcriptional activation of a complementary (co)-receptor subunit is necessary to reconstitute a functional sensory receptor. However, as we have shown in two cases that artificial co-receptor expression is insufficient to reconstitute function of “ectopic” tuning receptor activity, we suspect that levels of ectopic tuning receptor might also need to be enhanced, and there are possibly other requirements for functionality, such as perireceptor proteins or morphological specializations of the neuron/sensillum.

Our atlas of PCD-blocked antennae allowed us to characterize the development of many lineages fated to die. These lineages exhibit diverse properties: the undead neurons we identified were of different precursor types within diverse sensillar classes. Some of these robustly express a tuning receptor gene; here PCD can counteract promiscuous receptor expression, serving as a further regulatory layer to ensure precision in receptor patterning in the mature sensory system. Other undead neurons lack a detectable tuning receptor; however, our detection of co-receptor transcripts even in these “empty” neurons supports the idea that broad co-receptor expression extends to cells destined to die, thereby reflecting a more amenable evolutionary substrate for subsequent re-emergence of new neuron types from such dying lineages. Why some neurons express a receptor gene and other do not is an interesting open question. One possibility is that this property reflects their evolutionary age: lineages that more recently evolved a PCD fate might retain the gene regulatory network to permit receptor expression, while evolutionarily older dying lineages might have drifted in fate thus losing the capacity to express specific tuning receptors. Whatever the reason, the absence of “empty” neurons in the extant olfactory system implies that during emergence of a new sensory pathway from a dying lineage, changes in the gene regulatory network to turn off PCD and turn on a tuning receptor must be closely coordinated.

In this context, our identification of the first TFs (Mamo and Slp2) required for specification of PCD in OSN lineages provides an important entry-point into understanding how life/death fate decisions occur and evolve. The broader expression of both of these TFs beyond lineages fated to die emphasizes their context-dependent function, presumably because they are embedded within gene regulatory networks that influence survival, death and/or differentiation of sensory lineages. Further characterization of these, and other, TFs in dying lineages will help reveal whether and how they directly control pro-apoptotic genes expression, and why they promote PCD in some lineages but not others. Such knowledge will be key to understand how patterning of PCD can change during evolution to generate or remove individual sensory neuronal populations, and how this is coordinated with the selective expression of receptors.

While several lineages are entirely condemned to death (e.g., ac3 Ir75d-expressed neurons), we found, unexpectedly, that some dying neurons represent subsets of normal surviving lineages (e.g., Or65a/b/c neurons in at4). Such heterogeneity can be interpreted in different ways. The phenomenon might be an adaptive trait, for example, to limit the numbers of just one class of neurons within a specific sensillum type. However, the variation in Or65a/b/c population size across different *D. melanogaster* genotypes argues against this possibility, although we cannot exclude that such intraspecific phenotypic diversity arises from local adaptation of specific strains. Alternatively, heterogeneous PCD might reflect promiscuity in transcriptional specification of PCD resulting from overlap in gene regulatory networks of surviving and dying lineages, akin to the promiscuity observed in receptor expression. In this context, the observed heterogeneity in PCD specification might reflect a transitory evolutionary state of these pathways, such as the initial stages of loss of a sensory population. In extreme cases, promiscuous PCD might result in sensilla devoid of OSNs, as observed in rare instances (Nava Gonzales et al., 2021).

Taken together, our work both provides new understanding of the multilevel mechanisms that define the functional precision of the *Drosophila* olfactory system and highlights previously-overlooked variability at each of these levels that might provide a substrate for the molecular and cellular changes in these sensory pathways over evolutionary timescales. This study sets the stage for comparison of the antennal neuronal populations of phylogenetically diverse drosophilids and other insects to trace and understand the evolutionary changes in the olfactory system.

## Supporting information

FiguresS1-15

## Acknowledgements

We are very grateful for the invaluable assistance of the UNIL Flow Cytometry Facility and Lausanne Genomic Technologies Facility, and thank Gwénaëlle Bontonou and Roman Arguello for advice on HCR FISH. We acknowledge the Vienna *Drosophila* Resource Center, the Bloomington *Drosophila* Stock Centre (NIH P40OD018537) and the Developmental Studies Hybridoma Bank (NICHD of the NIH, University of Iowa) for reagents. We thank Nikos Konstantinides and members of the Benton laboratory for discussions and comments on the manuscript. Research in K.M.’s laboratory was supported by a National Institutes of Health award (R35GM133209). Research in R.B.’s laboratory is supported by the University of Lausanne, an ERC Advanced Grant (833548) and the Swiss National Science Foundation (310030_219185).

## Author contributions

J.M. conceived the project and performed most experiments and analyses, and supervised S.C., who performed histological experiments. A.S.B. performed electrophysiological experiments in at1. D.L. contributed to initial snRNA-seq sample preparation and analyses. P.C.C. generated the *Ir75d promoter-CD4:tdGFP* transgenic line. A.J. performed electrophysiological experiments in ac4, with input from and supervision by K.M. R.B. conceived and supervised the project. J.M. and R.B. wrote the paper with input from all co-authors.

## Methods

### *Drosophila* culture and transgenic line generation

Flies were reared in vials containing standard wheat flour/yeast/fruit juice medium and in incubators with 12 h light:12 h dark cycles at 25°C. Published strains are listed in Table S1.

The *Ir75d* promoter-CD4:tdGFP construct was generated by amplifying a 1994 bp DNA fragment from genomic DNA of the reference *D. melanogaster* strain (RRID:BDSC_2057) using the following forward and reverse PCR-primers GGGGACAAGTTTGTACAAAAAAGCAGGCTTCAgcaatggtaatattaaacta and GGGGACCACTTTGTACAAGAAAGCTGGGTCatccggcaactgattgcccca; this region encompasses 1850 bp 5’ regulatory sequence and 144 bp of exon 1, as in a previous promoter construct (Silbering et al., 2011). The amplified sequence was inserted into pDESTHemmarG (Addgene #31221) via Gateway recombination and confirmed by sequencing. The construct was integrated into attP40 (chromosome II) using phiC31-mediated transgenesis by BestGene Inc.

### Antennal dissection and nuclear isolation

10-15 virgin *peb-Gal4* females were placed in vials with 5-10 *UAS-unc84:GFP* or *UAS-p35,UAS-un84:GFP* males for 5 days, after which adults were removed. White pupae (corresponding to 0 h after puparium formation (APF)) were carefully transferred to fresh vials and aged for 18, 24, 30, 36, 42, 48, 56, 64, 72 or 80 additional hours. Developing antennae from aged pupae were dissected in ice cold Schneider’s medium (Gibco, 21720024) and immediately transferred to 1.8 ml Eppendorf tubes containing 100 µl Schneider’s medium, flash-frozen in liquid nitrogen and stored at -80°C. The numbers of antennae dissected in control (*peb-Gal4/+,UAS-unc84:GFP/+*) and PCD-blocked (*peb-Gal4/+,UAS-p35/+,UAS-unc84:GFP/+*) genotypes are as follows (time-point in h APF (n antennae control / n antennae PCD-blocked): 18 (49 / 57), 24 (72 / 52), 30 (67 / 56), 36 (52 / 51), 42 (45 /48), 48 (45 / 55), 56 (59 / 49), 64 (48 / 60), 72 (54 / 62) and 80 (47 / 53).

Samples were thawed on dry ice and nuclear suspensions prepared as described (Li et al., 2022; McLaughlin et al., 2021). Suspensions from “early” (18/24/30 h APF), “mid” (36/42/48 h APF) and “late” (56/64/72/80 h APF) developmental time points were pooled together, with the exception of time point 80 h APF in the control genotype, which was pooled with mid time points. This latter pooling reflected the initial experimental design, but 80 h APF control cells could be effectively re-classified in the late time point for all subsequent analyses. After addition of Hoechst 33342 (Thermo Fisher Scientific, 62249), samples were loaded into a FACSAria flow cytometer.

### Single-nuclear RNA-sequencing

For each pooled sample, 2 × 20,000 GFP-positive nuclei were sorted (20,000 for early nuclei in PCD-blocked condition), which were immediately loaded onto the Chromium Next GEM Chip (10x Genomics). Sequencing libraries were prepared with the Chromium Single Cell 3 reagent kit v3.1 dual index, following the manufacturer’s recommendations^LJ^. Libraries were quantified by a fluorometric method and quality was assessed on a Fragment Analyzer (Agilent Technologies). Sequencing was performed on an Illumina NovaSeq 6000 v1.5 flow cell for 100 cycles according to 10x Genomics recommendations (28 cycles read1, 10 cycles i7 index read, 10 cycles i5 index, and^LJ^91 cycles read2). Demultiplexing was performed with bcl2fastq2 Conversion Software (v2.20, Illumina). Raw snRNA-seq data was first processed through Cell Ranger (v6.1.2, 10x Genomics) with default parameters except -include introns-that was set to TRUE. A custom *D. melanogaster* reference genome and transcriptome from FlyBase (Drosophila_melanogaster.BDGP6.28.101) were used for mapping. Two marker genes, *GFP* and *p35*, were added to the GTF and FASTA files prior to building the custom genome reference with cellranger mkref (v6.1.2) function, following the 10x Genomics protocol.

### Integration of developing antennal snRNA-seq datasets from control and PCD-blocked genotypes

Ambient RNA contamination removal was applied on each of the cellranger output matrices from early, mid and late pools in control and PCD-blocked genotypes using SoupX (Young and Behjati, 2020) (v1.6.1, default parameters) and then subsequently integrated and analyzed using Seurat (v4.3.0.1) in RStudio. Matrices were normalized using SCTransform normalization (Hafemeister and Satija, 2019) (default parameters) and integrated using reciprocal PCA workflow (SelectIntegrationFeatures (nfeatures = 3000), PrepSCTIntegration, FindIntegrationAnchors (reference=control) and IntegrateData) described in (https://satijalab.org/seurat/articles/integration_rpca.html). PCA was used for clustering of the integrated datasets as follows: RunPCA (npcs=50), RunUMAP (reduction=”pca”, dims=1:50), FindNeighbors(reduction=”pca”, dims=1:50), FindClusters (resolution=0.5), resulting in 49 clusters.

### Cell type annotation

Marker genes (cutoff used: log_2_FC>3) of various cell types composing the adult antenna (sensory neurons, epithelial cells, hemocytes, muscle cells, glial cells and Johnston’s organ cells) were extracted from the Fly Cell Atlas dataset (Li et al., 2022) via the SCope interface (Davie et al., 2018). For support cells, *cut* and *shaven* were used as marker genes. For each cluster, the expression level (score) of cell type marker genes was computed using the AddModuleScore function implemented in Seurat. Cells were annotated through manual inspection of cell type scores in each cluster.

### Neuron class annotation

OSNs and other antennal sensory neurons from the control and PCD-blocked integrated datasets were subclustered at high resolution as follows RunPCA (npcs=45), RunUMAP (reduction=”pca”, dims=1:45), FindNeighbors(reduction=”pca”, dims=1:45, k.param=10), FindClusters (resolution=6), resulting in 162 clusters. The expression of diagnostic chemosensory receptor genes in the control dataset was used for initial cluster annotation. Clusters expressing more than one diagnostic receptor were further subclustered and annotated following the same pipeline. Because chemosensory receptor gene expression occurs relatively late during antennal development, these genes could not be used alone to discriminate neuron classes at earlier developmental stages. We therefore iteratively extracted marker genes of each neuron class as follows: FindAllMarkers (assay = “SCT”, logfc.threshold = 0.25, min.pct = 0.25, only.pos = T, test.use = “wilcox”, p_val_adj < 0.05) and evaluated the expression level of these markers in the unannotated clusters using the AddModuleScore function. Unannotated clusters showing the highest scores for a given lineage were assigned to that lineage, which were then incorporated into the next iteration of marker extraction and scoring. In cases of conflicting scoring of various marker lists, we manually inspected the expression of few top marker genes, and privileged shortest and continuous differentiation trajectories of lineages. After three iterations (0,1,2) of cluster annotation based on individual neuron classes, further annotation of the remaining unannotated clusters was based on sensillum type by grouping co-housed OSN lineages to extract sensillar markers as follows: FindAllMarkers (assay = “SCT”, logfc.threshold = 0.25, min.pct = 0.25, only.pos = T, test.use = “wilcox”, p_val_adj < 0.05) followed by AddModuleScore of the various marker genes. Overall, this iterative process allowed us to annotate 90% of the neurons; the remaining unannotated fraction mostly correspond to early phase cells. Marker genes distinguishing coeloconic and sacculus neuron populations were extracted from (Corthals et al., 2023; Mika et al., 2021). Gene expression levels shown in the dot plots and UMAPs are residuals from a regularized negative binomial regression, and have arbitrary units.

### Lineage developmental and pseudotime reconstruction

Individual sensillar lineages were extracted from the integrated datasets and subclustered using a similar workflow as described above but with lineage-specific parameters: RunPCA, RunUMAP, FindNeighbors, FindClusters. Each cluster was assigned to early, mid and late developmental phases based on PCD-blocked data and then extrapolated to the control data (this was necessary, due to the pooling of the 80 h APF time point with the mid stages in the latter). Individual lineages were imported in monocle3 (v1.2.9) (Cao et al., 2019; Levine et al., 2015; Qiu et al., 2017; Trapnell et al., 2014) and pseudotime inferred as follows: cluster_cells, learn_graph (use_partition = F), order_cells (start_end was chosen at the tip of “early” stage cells) and gene expression dynamics plotted using plot_genes_in_pseudotime (color_cells_by = “pseudotime”) function.

### Identification of undead neurons

During reconstruction of individual lineages, we systematically checked for the presence of clusters that would be exclusive to the PCD-blocked dataset using DimPlot (split.by = “condition”) function in Seurat. We confirmed that any such PCD-blocked specific clusters corresponded to undead neuron lineages by quantifying the expression of the pro-apoptotic genes *rpr*, *grim* and *skl* (RGS score) using AddModuleScore function. If there was no difference in clusters between control and PCD-blocked animals, we quantified the number of cells (from 36 h APF) of each lineage in both conditions. In a few cases (e.g., at4 Or65a/b/c neurons) there was a clear increase in number in the PCD-blocked dataset. We further validated such additional cells as being undead neurons, by quantifying and ranking RGS score at the sensillar level using AddModuleScore, ViolinPlot (features = “RGS”, sort = T) function. Typically, lineages with “embedded” undead neurons had the highest RGS score.

### Differential gene expression analysis

Differentially expressed genes (DEGs) were extracted from various Seurat objects using FindAllMarkers function with the following parameters applied for various comparisons: for developmental time marker genes (Figure S2): (object = “All OSNs”, assay = “SCT”, logfc.threshold = 0.25, min.pct = 0.25, only.pos = T, test.use = “wilcox”, p_val_adj < 0.05), for OSN marker genes (Figure S5): (object = “All annotated OSNs in control condition”, assay = “SCT”, logfc.threshold = 0.25, min.pct = 0.18, only.pos = T, test.use = “wilcox”, p_val_adj < 0.05), for sensilla marker genes (Figure S6): (object = “All annotated OSNs in control condition”, assay = “SCT”, logfc.threshold = 0.25, min.pct = 0.25, only.pos = T, test.use = “wilcox”, p_val_adj < 0.05), for OSN precursor type marker genes (Figure S7): (object = “All annotated OSNs in control condition except arista”, assay = “SCT”, logfc.threshold = 0.25, min.pct = 0.4, only.pos = T, test.use = “wilcox”, p_val_adj < 0.05). For each comparison, an analysis of Gene Ontology terms over-represented in marker genes was performed using clusterProfiler (v4.4.4) (Wu et al., 2021).

### Hierarchical clustering of sensory neurons based on differentially expressed transcription factors

A phylogenetic tree relating sensory neuron classes (Figure 3H) was built based on a distance matrix constructed in the differentially expressed transcription factors (DE_TFs) space (Figure S8) using the BuildClusterTree function implemented in Seurat: BuildClusterTree (assay = “SCT”, features = DE_TFs).

### Hybridization chain reaction RNA fluorescent *in situ*

RNA probes (Table S2) were synthesized by Molecular Instruments, and we followed a published HCR RNA FISH protocol (Bontonou et al., 2024) with minor modifications. Female flies (2-7 days old) of interest were flash-frozen in liquid nitrogen and antennae were passed through a mini-sieve (mesh-width = 80 µm) and collected in Petri dishes containing fixation solution (1×PBS, 3% Triton X-100, 4% paraformaldehyde). For each staining, 30-40 antennae were immediately transferred into 1.5 ml Eppendorf tubes filled with 1 ml of fixation solution and place on a platform for 1 h at room temperature (RT) with gentle shaking (35 rpm). Fixative solution was removed and samples washed 2× 5 min with 1 ml of PBS 1×, 3% Triton X-100 at RT, followed by 3× 5 min with 1 ml of 1×PBS, 0.1% Triton X-100 at RT. Wash solution was removed and samples pre-hybridized with 300 µl of probe hybridization buffer (Molecular Instruments) for 30 min at 37°C. Pre-hybridization buffer was removed and replaced by probe solution (5 µl of probe in hybridization solution, 500 µl final volume, except for *Ir75d*, *Ir41a*, *Or85f*, *Or88a, Or67d* and *Ir76a* probes for which 10 µl were used) for 16 h at 37°C. Probe solution was removed and samples washed 4× 15 min with 500 µl of pre-heated (37°C) probe wash buffer (Molecular Instruments) at 37°C, followed by 3× washes of 5 min at RT with 500 µl of 5×SSCT solution (5×SSC, 0.1% Triton X-100). Samples were pre-amplified with 300 µl of amplification buffer (Molecular Instruments) for 30 min at RT. During this time, for each probe in a sample, 10 µl of hairpin amplifiers h1 and 10 µl of hairpin amplifiers h2 (or 15 µl each when probe volume was doubled) were heated at 95°C for 90 s, and snap-cooled to RT, with protection from light. Pre-amplification buffer was removed from the samples and replaced by the snap-cooled amplifiers in amplification buffer (Molecular Instruments), 500 µl final volume, and placed at RT and protected from light for 16 h. Amplification solution was removed and samples washed 5× in 500 µl of 5×SSCT at RT (2× 5 min, 2× 30 min and 1× 5 min). 5×SSCT was removed and 50 µl of Vectashield mounting medium (Vector Laboratories, H-1200) was added before mounting.

### Standard RNA fluorescent *in situ* hybridization and immunohistochemistry

For the experiments in Figure 6G, we used a standard RNA FISH protocol (Saina and Benton, 2013) on female flies (2-7 day old), using RNA FISH probes generated using primers in Table S2. When combined with immunohistochemistry (Figure S15A), we performed standard RNA FISH until TSA-Cy5 colorimetric detection, when samples were washed 5× 20 min at RT with 500 µl of TNT buffer and incubated for 1 h in 300 µl of blocking solution (1×PBS, 0.2% Triton X-100, 5% heat inactivated goat serum). Supernatant was replaced by 500 µl of anti-GFP-chicken antibody containing blocking solution and samples placed at 4°C on a rotating wheel for 40 h. Samples were then washed 5× 20 min at RT with 500 µl of 1×PBS, 0.2% Triton X-100, and incubated for 1 h in 300 µl of blocking solution. Supernatant was replaced by 500 µl of secondary antibodies in blocking solution and samples placed at 4°C on a rotating wheel for 40 h. Samples were washed 5× 20 min at RT with 500 µl of 1×PBS, 0.2% Triton X-100 and 50 µl of Vectashield mounting medium (Vector Laboratories, H-1200) was added to each sample prior to mounting. Primary and secondary antibodies are listed in Table S3.

### Immunohistochemistry in central brain

Female flies (2-7 day old) were fixed in 2 ml of PB 1×, 3% Triton X-100, 4% paraformaldehyde for 2 h at RT. Brains were dissected into ice-cold 1×PB (under a binocular microscope and immediately transferred into 1.5 ml Eppendorf tubes containing 1 ml of 1×PB, 0.3% Triton X-100. Brains were washed 5× 15 min in 1 ml of 1×PB, 0.3% Triton X-100 at RT, and then incubated in 1 ml of blocking solution (PB 1×, 0.3% Triton X-100, 5% heat-inactivated goat serum) for 1 hat RT. Supernatant was replaced by 500 µl of primary antibodies in blocking solution and the samples placed on a rotating wheel for 40 h at 4°C. Brains were washed 5× 15 min in 1 ml of 1×PB, 0.3% Triton X-100 at RT, and then incubated in 1 ml of blocking solution for 1 h at RT. The supernatant was replaced by 500 µl of secondary antibodies containing blocking solution and the samples placed on a rotating wheel for 40 h at 4°C. Brains were washed 5× 15 min in 1 ml of 1×PB, 0.3% Triton X-100 at RT, and 50 µl of Vectashield mounting medium (Vector Laboratories, H-1200) was added prior to mounting.

### Image acquisition and processing

Images from antennae and antennal lobes were acquired with confocal microscopes (Zeiss LSM710 or Zeiss LSM880 systems) using a 40× (antennae) or a 63× (antennal lobe) oil immersion objective. Images were processed using Fiji software (Schindelin et al., 2012).

### Electrophysiology

Single-sensillum recordings on ac4 coeloconic sensilla were performed on 2-4 day old female flies using glass electrodes filled with sensillum recording solution, essentially as described (Vulpe et al., 2021). Coeloconic sensilla were identified based on their stereotyped locations on the antenna and responses to diagnostic odorants. The odor response was calculated from the difference in summed spike frequencies of all OSNs in response to a 0.5 s odor puff compared to a 0.5 s solvent puff, as described (Vulpe et al., 2021).

Single sensillum recordings from at1 were performed on 5-10-day old female flies using tungsten electrodes, essentially as described (Benton and Dahanukar, 2023); sensilla were identified by their morphology, characteristic location and responses of Or67d neuron to cVA. Odor responses were calculated from the difference between the OSN spike frequency during and before odor stimulation, as described (Benton et al., 2007), from which the solvent response was subtracted. Spike amplitudes numbers in at1 (Figure 7D) were scored independently by two experimenters blind to the genotype. Information on odor stimuli and the paraffin oil solvent is provided in Table S4.

## Supplementary Information

**Data S1. Odor-evoked neuronal responses.**

*- see separate Excel file*

**Data S2. Top marker genes in undead and normal Ir75d neurons.**

*- see separate Excel file*

**Data S3. RNAi screen data.**

*- see separate Excel file*

**Table S1.**
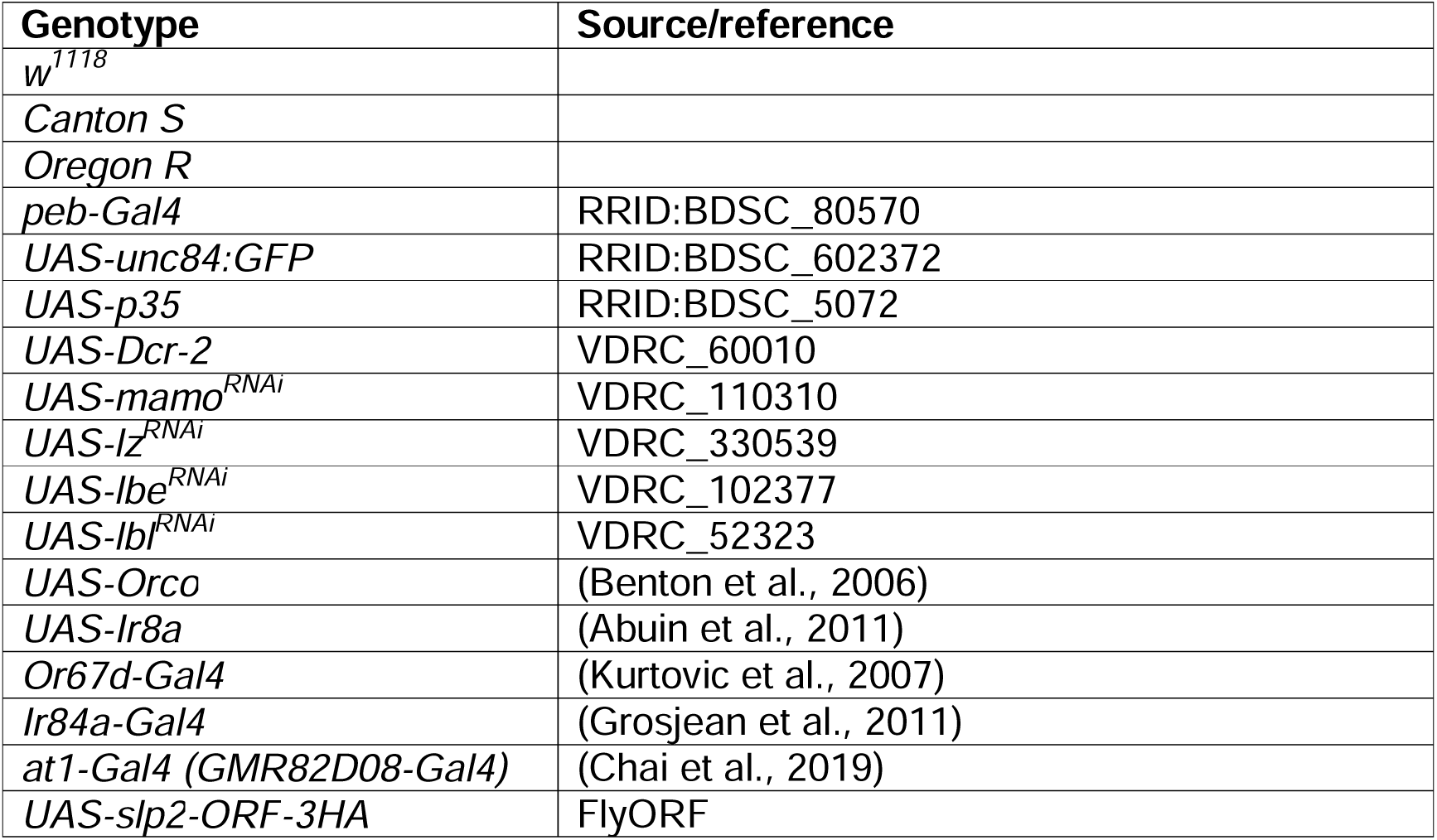
*Drosophila* strains.

**Table S2.**
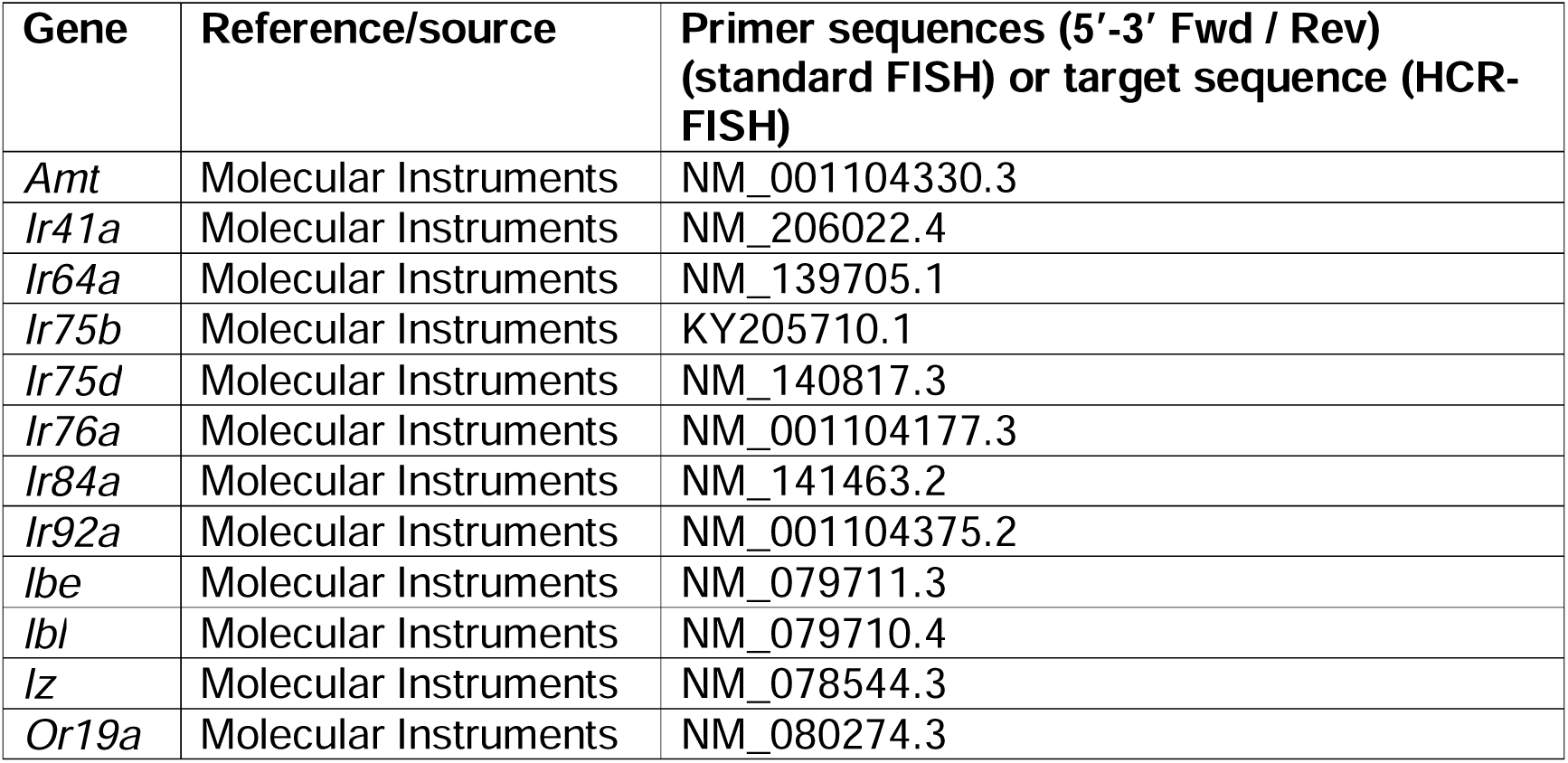
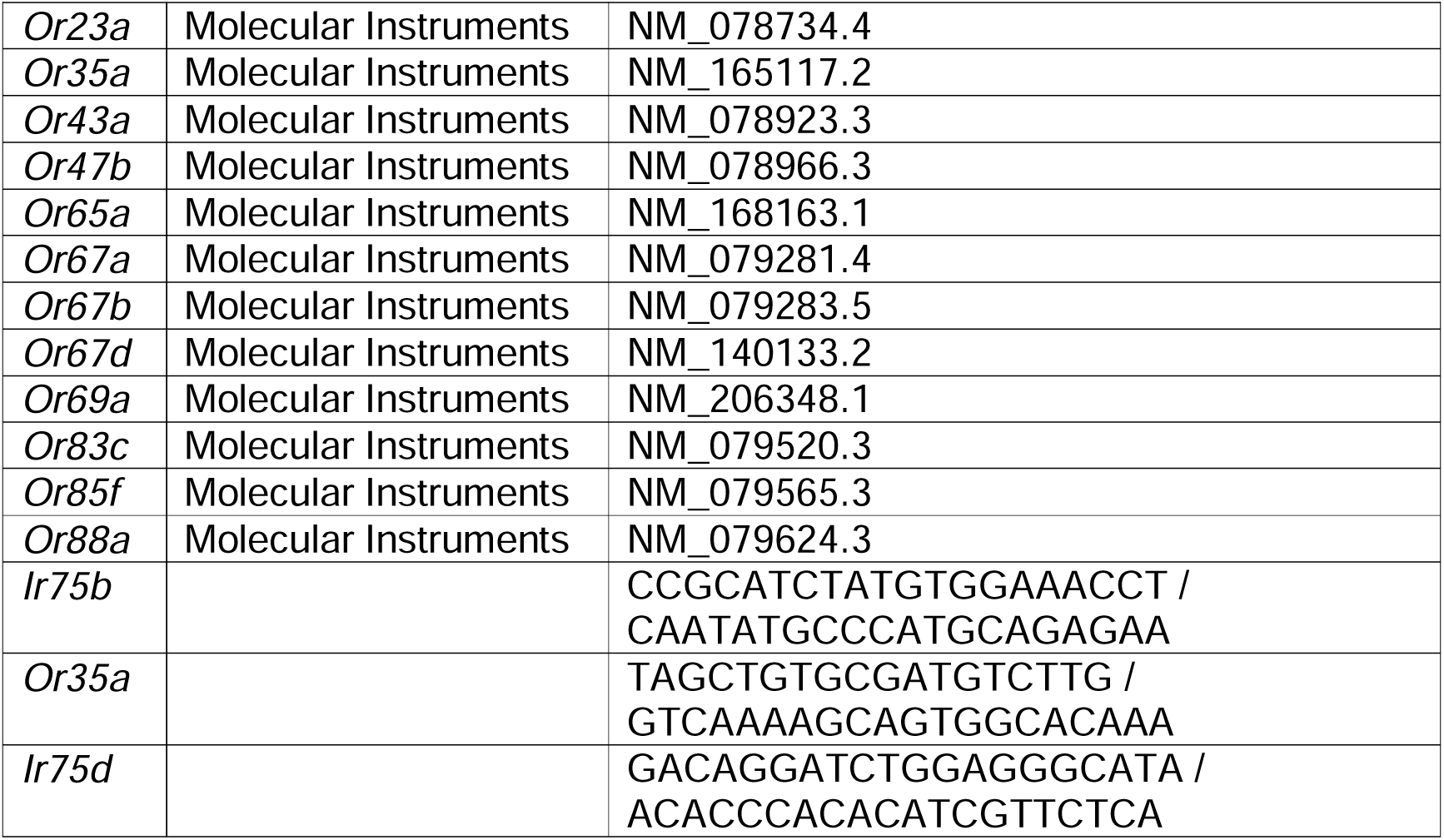
RNA FISH probes.

**Table S3.** Antibodies.

| **Antibody** | **Dilution** | **Reference/source** | **Identifier** |
| --- | --- | --- | --- |
| Chicken anti-GFP | 1:500 | Abcam | ab13970 |
| Mouse anti-Bruchpilot (nc82) | 1:10 | DSHB | nc82 |
| anti-DIG-POD | 1:300 | Roche Diagnostics AG | 11 207 733 910 |
| anti-Fluorescein-POD | 1:300 | Roche Diagnostics AG | 11 426 346 910 |
| Alexa Fluor 488 Goat anti-Chicken | 1:500 | Abcam | ab150169 |
| Cy5 Goat anti-Mouse | 1:250 | Jackson Immunoresearch | 115-175-166 |

**Table S4.** Odors.

| **Odor** | **CAS** | **Source** |
| --- | --- | --- |
| *11-cis*-vaccenyl acetate (cVA) | 6186-98-7 | Pherobank |
| *E2*-hexenal | 6728-26-3 | Sigma-Aldrich |
| 1-hexanol | 111-27-3 | Acros Organics |
| hexyl acetate | 142-92-7 | Sigma-Aldrich |
| 1-octanol | 111-87-5 | Acros Organics |
| 2-oxopentanoic acid | 1821-02-9 | Sigma-Aldrich |
| paraffin oil (solvent) | 8012-95-1 | Acros Organics |
| phenethylamine | 64-04-0 | Acros Organics |
| phenylacetaldehyde | 122-78-1 | Alfa Aesar |

## Supplementary Figures

**Figure S1. Global cell type annotation of the developing antenna.**

**(A)** Distribution of the number of genes detected per nucleus across all sequenced nuclei in control and PCD-blocked datasets. The dashed vertical lines indicate the mean.

**(B)** UMAPs illustrating the unsupervised clustering of all nuclei from the integrated control and PCD-blocked datasets (top left), cell type scoring using marker gene modules extracted from the Fly Cell Atlas (Li et al., 2022) (see Methods), and cell type annotation of the integrated datasets (bottom right). Nuclei from clusters 15 and 26 were sparse and exhibited a mixed identity; we therefore assigned these as doublets and discarded them from downstream analyses.

**(C)** Expression score of cell type marker gene modules and the fraction of detected mitochondrial genes in each cluster.

**(D)** Cell type and nuclei number composing the integrated, control and PCD-blocked datasets.

**Figure S2. Hallmarks of peripheral olfactory system development.**

**(A)** Expression score of early, mid and late developmental time marker gene modules in early, mid and late olfactory, hygrosensory and thermosensory neurons, integrated datasets. Boxes show the median (thick line), first and third quartiles, while whiskers indicate data distribution limits. A-C letters indicate significant differences: P < 0.05 in pairwise comparisons (Wilcoxon rank sum test followed by Bonferroni correction for multiple comparisons).

**(B)** Expression of the top 30 genes (log_2_FC) of each developmental time marker module in neurons grouped by developmental time (integrated datasets).

**(C)** Gene Ontology (GO) analysis illustrating the top 10 (log_10_(adjusted P)) Biological Process (BP), Molecular Function (MF) and Cellular Component (CC) categories enriched in early, mid and late developmental marker gene modules (810 genes total).

**Figure S3. Initial annotation of neuronal subclusters based upon sensory receptor expression.**

**(A)** UMAPs of unsupervised clustering of the sensory neurons (integrated datasets) at iteration 0 (left) and the initial annotation of a subset of these (typically late-stage cells) based upon sensory receptor gene expression in the control dataset (right).

**(B)** Expression of diagnostic *Or* and *Ir* genes (and the glial marker *repo*) in each cluster from (A) (left UMAP, control dataset only).

**(C)** UMAPs of unsupervised sub-clustering of the multiple_OR cluster from (A) (right UMAP) (top) and the annotation based upon sensory receptor gene expression in the control dataset (bottom).

**(D)** Expression of diagnostic *Or* and *Ir* genes in each cluster in (C) (top UMAP; control dataset only).

**Figure S4. Backward, iterative annotation of subclusters based upon sensory receptor expression.**

**(A)** UMAPs at iteration 0 of sensory neurons annotation (left) and highlighting at4 neurons Or88a (red), Or65a/b/c (green) and Or47b (blue) (right).

**(B)** Ranked expression score (left-to-right) of each at4 OSN marker gene module extracted from the iteration 0 annotated dataset. The top 5 unannotated clusters with the highest scores for each OSN population marker genes modules are shown, allowing us to assign clusters from earlier developmental stage (typically lacking sensory receptor expression) to each at4 OSN, as indicated in the text on the right. These clusters were used for a subsequent round of marker gene scoring.

**(C)** UMAPs at iteration 1 of sensory neurons annotation (left) and highlighting Or88a (red), Or65a/b/c (green) and Or47b (blue) neurons (right).

**(D)** Ranked expression score (left-to-right) of each at4 OSN marker gene module extracted from the iteration 1 annotated object. The top 5 unannotated clusters with the highest scores for each OSN population marker genes modules are shown, allowing us to assign clusters to Or88a but not Or47b or Or65a/b/c OSNs, as indicated in the text on the right). These clusters were used for a subsequent round of marker gene scoring.

**(E)** UMAPs at iteration 2 of sensory neurons (left) and highlighting Or88a (red), Or65/a/b/c (green) and Or47b (blue) neurons (right). No further sensory neuron lineage-based backward annotation was possible.

**(F)** UMAP at iteration 2 (as in (E)) with sensilla-based annotations.

**(G)** Ranked expression score (left-to-right) of the at4 sensillum marker gene module extracted from the iteration 2, sensilla based, annotated object. The top 5 unannotated clusters with the highest scores are shown, allowing us to assign clusters to at4, as indicated in the text on the right. These clusters were used for a subsequent round of marker gene scoring.

**(H)** UMAPs at iteration 3 of the sensilla-based annotation of sensory neurons (left) and highlighting the at4 sensillum (right).

**Figure S5. Sensory neuron type marker genes.**

**(A)** GO analysis illustrating the top 25 (log_10_(adjusted P)) Biological Process (BP), Molecular Function (MF) and Cellular Component (CC) categories enriched in OSNs marker gene modules (1242 genes total). Only the control dataset was analyzed in this and the following panel.

**(B)** Expression of OSN marker genes belonging to the indicated GO categories.

**Figure S6. Sensilla marker genes.**

**(A)** Expression score of sensilla marker gene modules across sensilla (control dataset only in this and other panels).

**(B)** Gene Ontology (GO) analysis illustrating the top 25 (log_10_(adjusted P)) Biological Process (BP), Molecular Function (MF) and Cellular Component (CC) categories enriched in sensilla marker gene modules (727 genes total).

**(C)** Expression of the top 5 marker genes (log_2_FC) of each sensillum class.

**Figure S7. Sensory neuron precursor type marker genes.**

**(A)** Expression score of Naa, Nab, Nba and Nbb marker gene modules in each sensory neuron precursor type category. Boxes show the median (thick line), first and third quartiles, while whiskers indicate data distribution limits. A-D letters indicate significant differences: P < 0.05 in pairwise comparisons (Wilcoxon rank sum test followed by Bonferroni correction for multiple comparisons). Only the control dataset was analyzed in this and the following panels.

**(B)** Expression of the top 20 genes (log_2_FC) from each of the sensory neuron precursor type marker gene modules across sensory neuron precursor types.

**(C)** Gene Ontology (GO) analysis showing the top 25 (log_10_(adjusted P)) Biological Process (BP), Molecular Function (MF) and Cellular Component (CC) categories enriched in OSN type marker gene modules (200 genes total).

**(D-E)** UMAPs of ac1 (D) and at4 (E) with annotation and developmental phases (left), the expression of diagnostic sensory receptors (middle), the expression score of sensory neuron precursor type marker gene modules (right) and a schematic illustrating the inferred precursor type and identity of sensory neurons (bottom left). A-D letters indicate significant differences: P < 0.05 in pairwise comparisons (Wilcoxon rank sum test followed by Bonferroni correction for multiple comparisons).

**Figure S8. Putative transcription factor codes underlying sensory neuron identity.**

Differentially expressed TFs in sensory neuron populations (control dataset). A manually curated TF reference list was extracted from FlyMine database (https://www.flymine.org/flymine). TFs functionally characterized in this work are highlighted.

**Figure S9. Sensory receptor expression during antennal development.**

Expression of tuning and co-receptor subunits across cell types from different developmental phases (control dataset).

**Figure S10. Examples of receptor co-expression.**

**(A)** Top: UMAPs of the sacIII_v/d-Amt lineage (control dataset) illustrating the developmental stages and receptor expression patterns; *Rh50* encodes an ammonia transporter that is co-expressed with Amt in these ammonia-sensing neurons although its role is unclear (Vulpe et al., 2021). Bottom: a pseudotime UMAP and corresponding receptor expression dynamics.

**(B)** Left: RNA FISH on whole-mount antennae of control (*peb-Gal4*) animals with probes targeting the indicated transcripts (*n* = 10-12). The ac3 and ac1 sensilla zones are indicated. Bottom row shows a higher magnification of sacIII-v/d-Amt neurons co-expressing *Amt* and *Or35a* in 3 adjacent confocal Z-slices. Scale bars, 25 µm (top images) or 10 µm (bottom images). Right: schematic of olfactory (co)-receptor subunits expressed in sacIII-v/d-Amt neurons.

**(C-E)** Top: UMAPs of the ac1-Ir31a (C), ab5B-Or47a/Or33b (D) and ab8B-Or9a (E) lineages (control dataset) illustrating the developmental stages and receptor expression patterns. Bottom: pseudotime UMAPs and corresponding receptor expression dynamics.

**Figure S11. Global survey of chemosensory receptor expression in late stages.** Expression of all detected sensory receptor subunits in the late developmental phase (control dataset).

**Figure S12. Expression of pro-apoptotic genes.**

**(A)** UMAPs illustrating the expression of the pro-apoptotic genes *reaper* (*rpr*), *grim*, *sickle* (*skl*) and *head involution defective* (*hid)* genes in annotated nuclei of control and PCD-blocked datasets.

**(B)** Average expression (left) and fraction of positive nuclei (right) for these genes in control and PCD-blocked datasets.

**(C)** Abundance of each neuronal class in control (top) and PCD-blocked (middle) datasets, calculated as the percentage of nuclei for each class relative to the total number of nuclei (from 36 h APF, excluding those forming new clusters in the PCD-blocked dataset) and fold-change in the abundance of each class in the PCD-blocked dataset relative to the control dataset (bottom).

**Figure S13. Expression of co-receptors in undead neurons.**

**(A)** UMAPs of the integrated ac3I/II dataset illustrating the expression of tuning receptors (as in Figure 5D) and co-receptors in endogenous and undead neurons.

**(B)** UMAPs of the integrated sacIII-v/d dataset illustrating the expression of tuning receptors (as in Figure 5K) and co-receptors in endogenous and undead neurons.

**Figure S14. Analysis of the undead neuron lineages in additional sensilla.**

**(A)** UMAPs of the ab10 lineage from control and PCD-blocked datasets illustrating the developmental stages (top) and RGS expression (bottom), revealing that some cells with high RGS score are present exclusively in PCD-blocked animals.

**(B)** Lineage annotation of the ab10 sensillum in control and PCD-blocked integrated datasets (top), ranked RGS score (left-to-right) (bottom left) and precursor type score (bottom right) for each OSN type in the integrated datasets. Boxes show the median (thick line), first and third quartiles, while whiskers indicate data distribution limits, here and elsewhere.

**(C)** Expression of the indicated receptor in the integrated control and PCD-blocked datasets. No receptor was robustly detected in the undead neuron population.

**(D)** Schematic of the inferred states of the ab10 sensillum in control and PCD-blocked antennae.

**(E)** UMAPs of the ab5 lineage from control and PCD-blocked datasets illustrating the developmental stages (top) and RGS expression (bottom), revealing that some cells with high RGS score are present exclusively in PCD-blocked animals.

**(F)** Lineage annotation of the ab5 sensillum in control and PCD-blocked integrated datasets (top), ranked RGS score (left-to-right) (bottom left) and precursor type score (bottom right) for each OSN type in the integrated datasets.

**(G)** Expression of the indicated receptor in the integrated control and PCD-blocked datasets. No receptor was robustly detected in the undead neuron population.

**(H)** Schematic of the inferred states of the ab5 sensillum in control and PCD-blocked antennae.

**(I)** UMAPs of the sacI lineage from control and PCD-blocked datasets illustrating the developmental stages (top) and RGS expression (bottom), revealing that some cells with high RGS score are present exclusively in PCD-blocked animals.

**(J)** Lineage annotation of the sacI sensillum in control and PCD-blocked integrated datasets (top), ranked RGS score (left-to-right) (bottom left) and precursor type score (bottom right) for each OSN type in the integrated datasets.

**(K)** Expression of the indicated receptor in the integrated control and PCD-blocked datasets. No receptor was robustly detected in the undead neuron population.

**(L)** Schematic of the inferred states of the sacI sensillum in control and PCD-blocked antennae.

**(M-N)** UMAPs of the ai3 (E) and ab4 (F) lineages from control and PCD-blocked datasets illustrating the developmental stages (top) and RGS expression (bottom). No undead neurons were apparent.

**(B,F,J)** A-D letters indicate significant differences: P < 0.05 in pairwise comparisons (Wilcoxon rank sum test followed by Bonferroni correction for multiple comparisons).

**Figure S15. Analysis of Ir75d neuron projection patterns.**

**(A)** *Ir75b/c* RNA FISH and anti-GFP immunofluorescence on whole-mount antennae of control (*peb-Gal4/+;Ir75d-CD4-GFP/+*; top), *mamo^RNAi^* (*peb-Gal4,UAS-Dcr-2/+;UAS-mamo^RNAi^/Ir75d-CD4-GFP;* middle) and PCD-blocked (*peb-Gal4/+;UAS-p35/Ir75d-CD4-GFP;* bottom) animals (*n* = 10, 11 and 6, respectively). Scale bars, 25µm.

**(B)** GFP and nc82 immunofluorescence on whole-mount brains of control (*peb-Gal4/+;Ir75d-CD4-GFP/+;* top), *mamo^RNAi^* (*peb-Gal4,UAS-Dcr-2/+;UAS-mamo^RNAi^/Ir75d-CD4-GFP*; middle) and PCD-blocked (*peb-Gal4/+;UAS-p35/Ir75d-CD4-GFP;* bottom) animals (*n* = 4, 8 and 6, respectively). Scale bars, 25 µm.

