## Supplementary material for "Multilayer regulation underlies the functional precision and evolvability of the olfactory system": FiguresS1-15

Mermet Figure S1

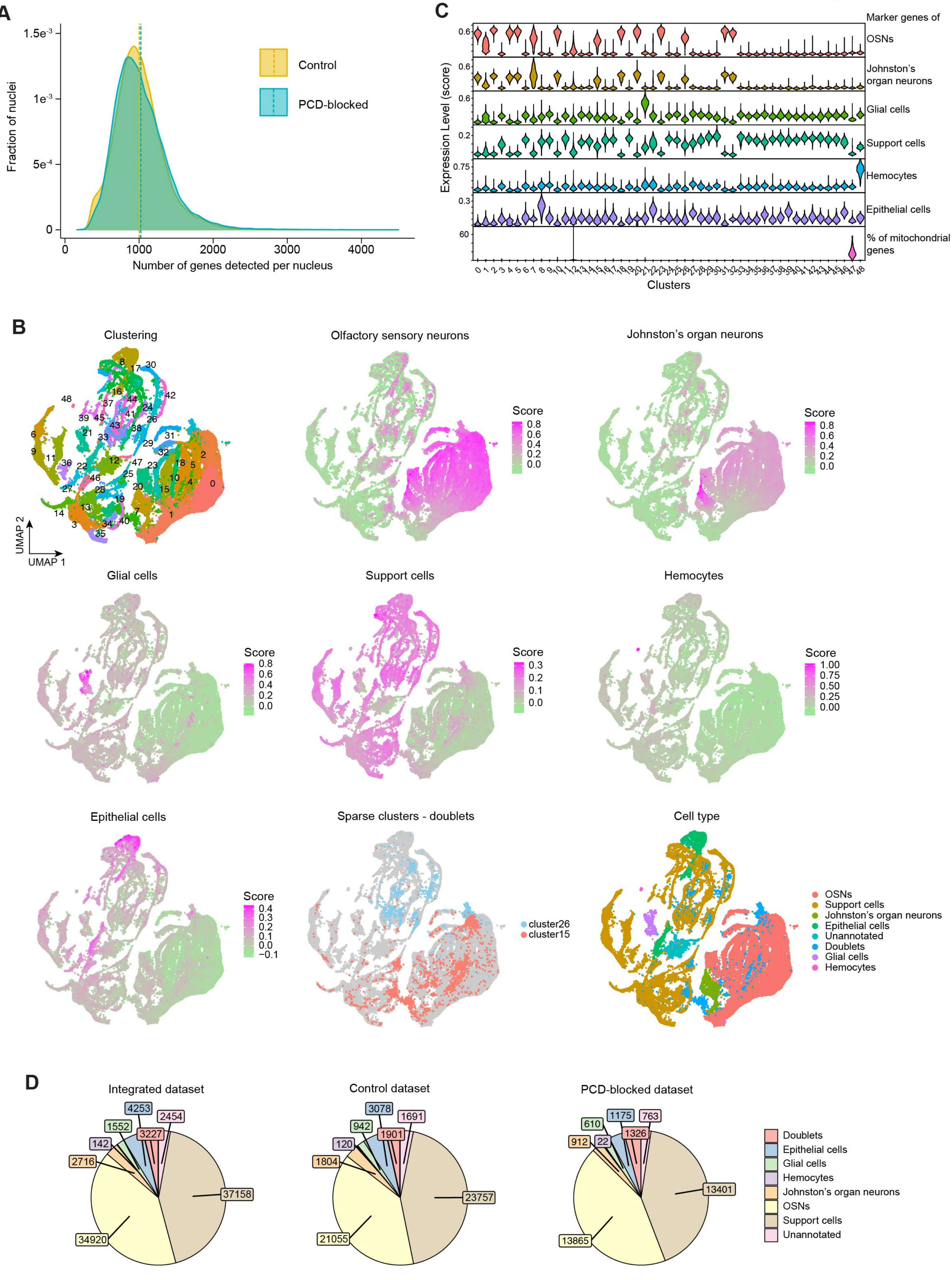

Mermet Figure S2

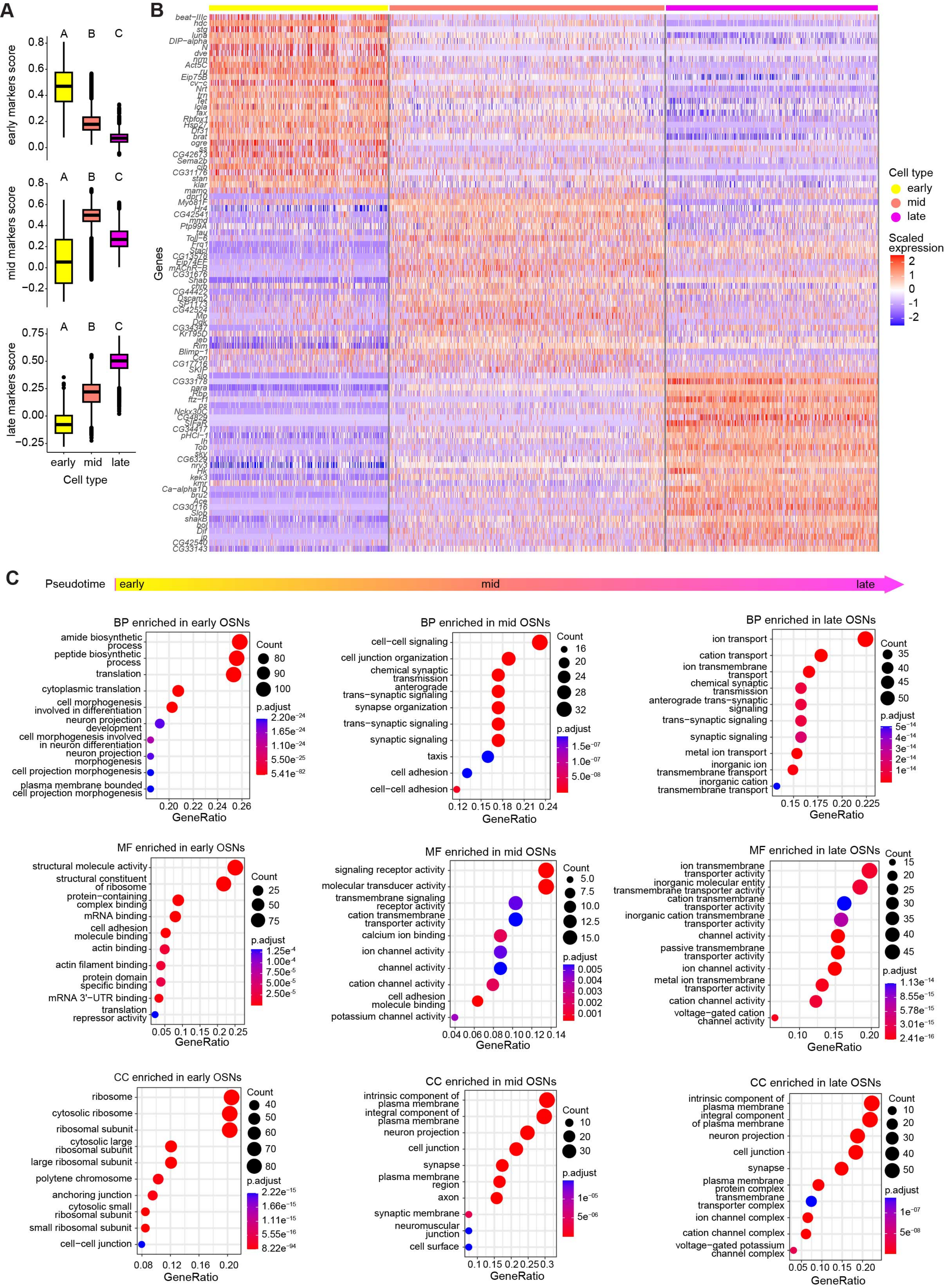

Mermet Figure S3

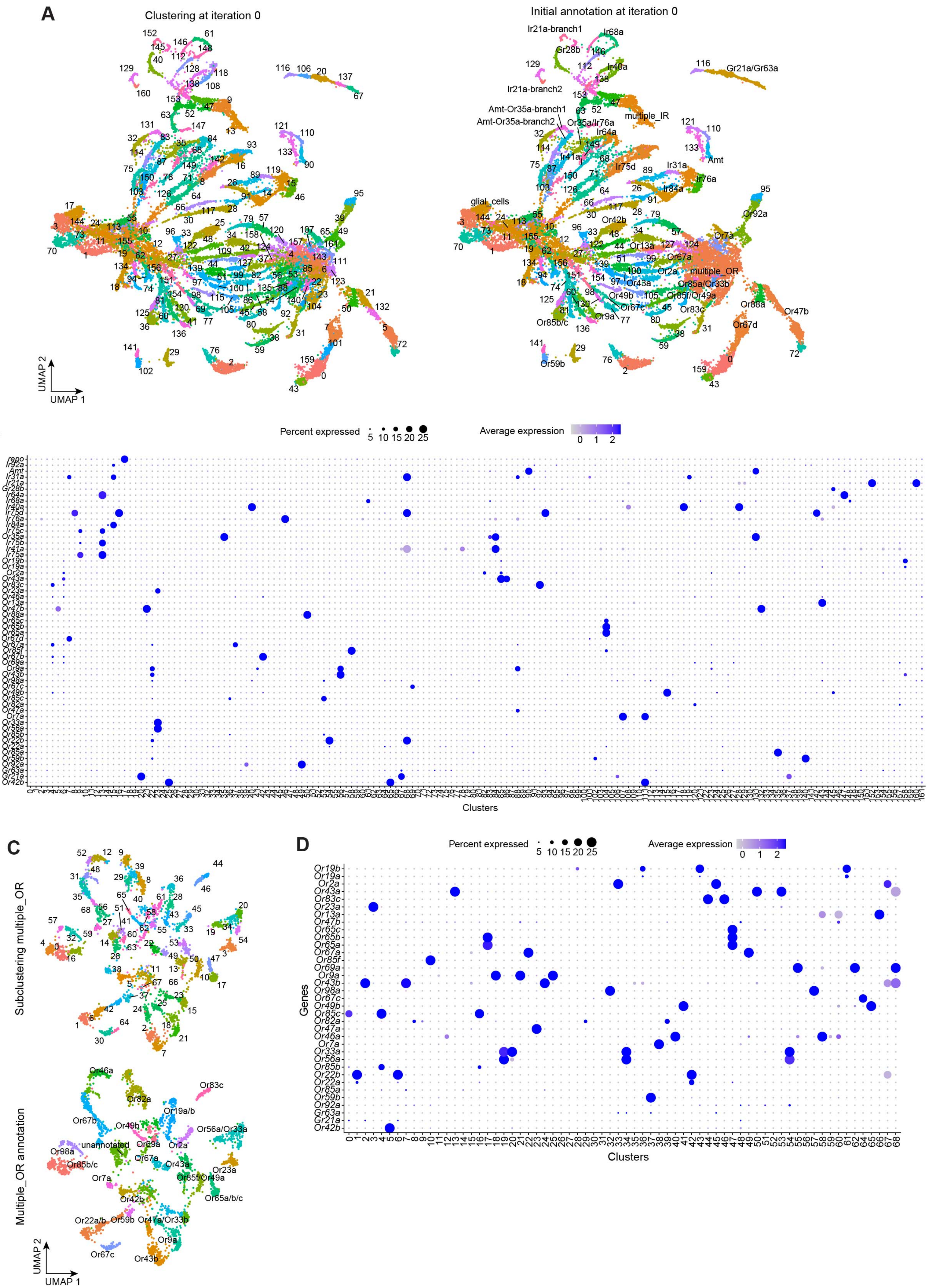

Mermet Figure S4

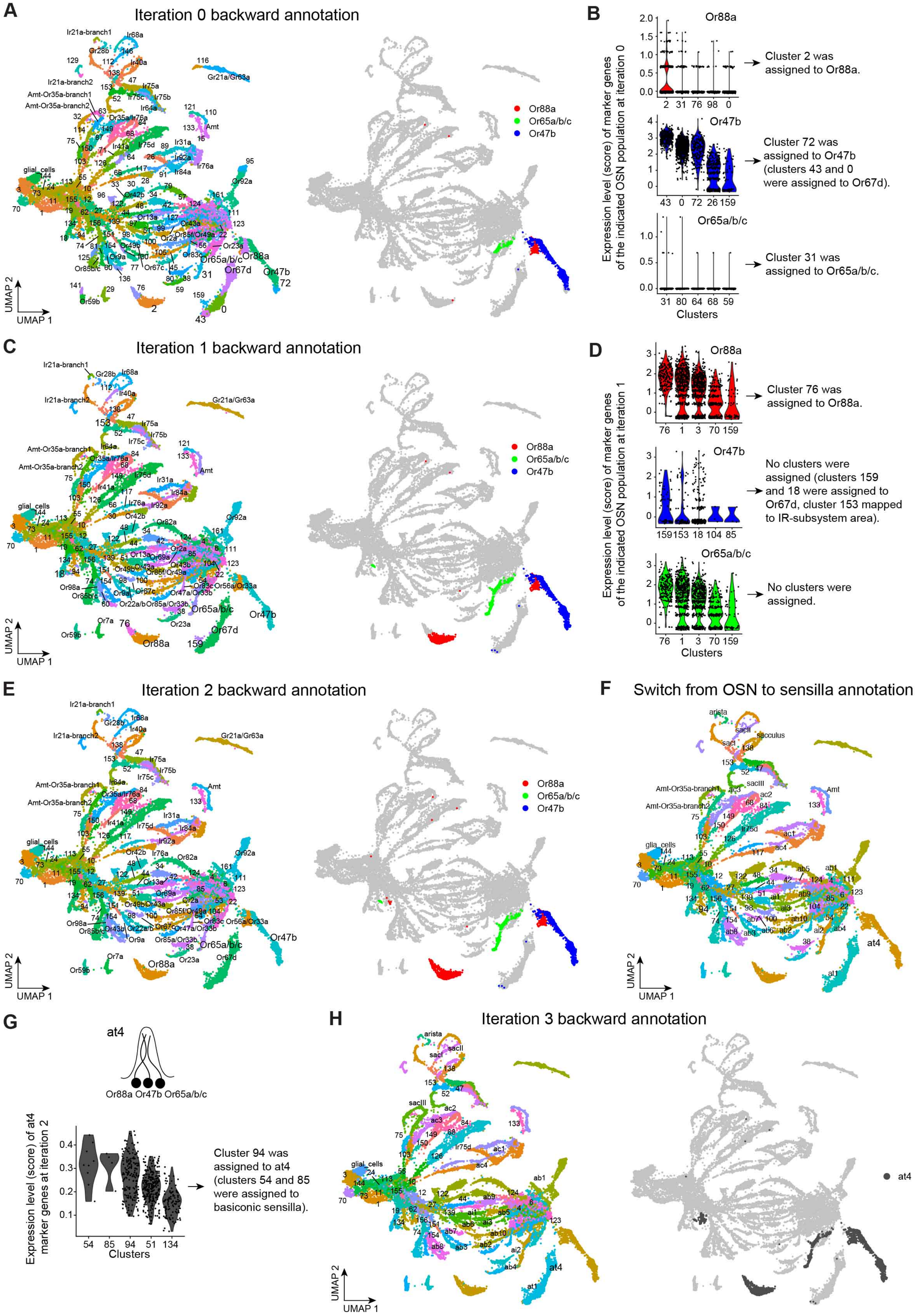

Mermet Figure S5

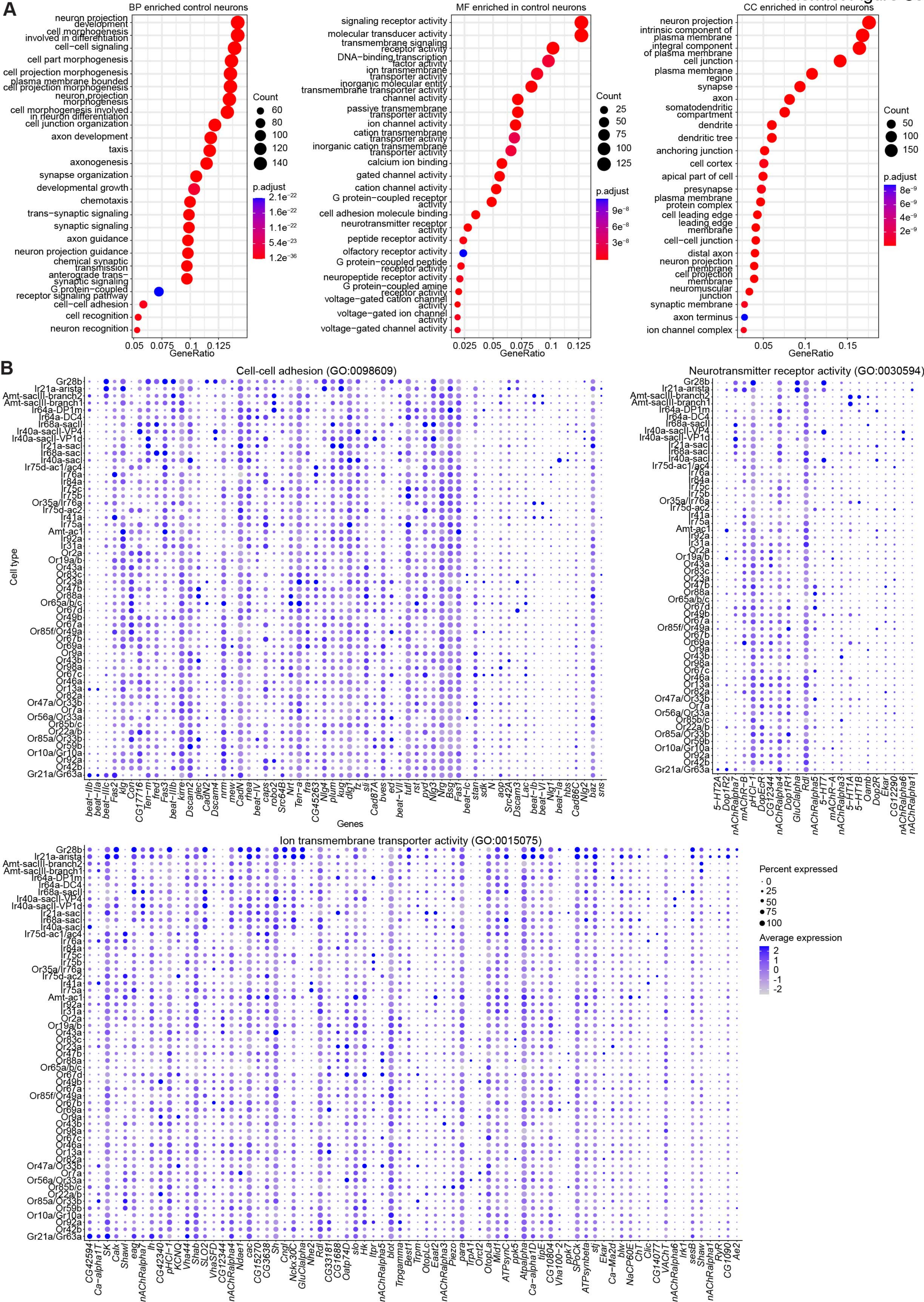

Mermet Figure S6

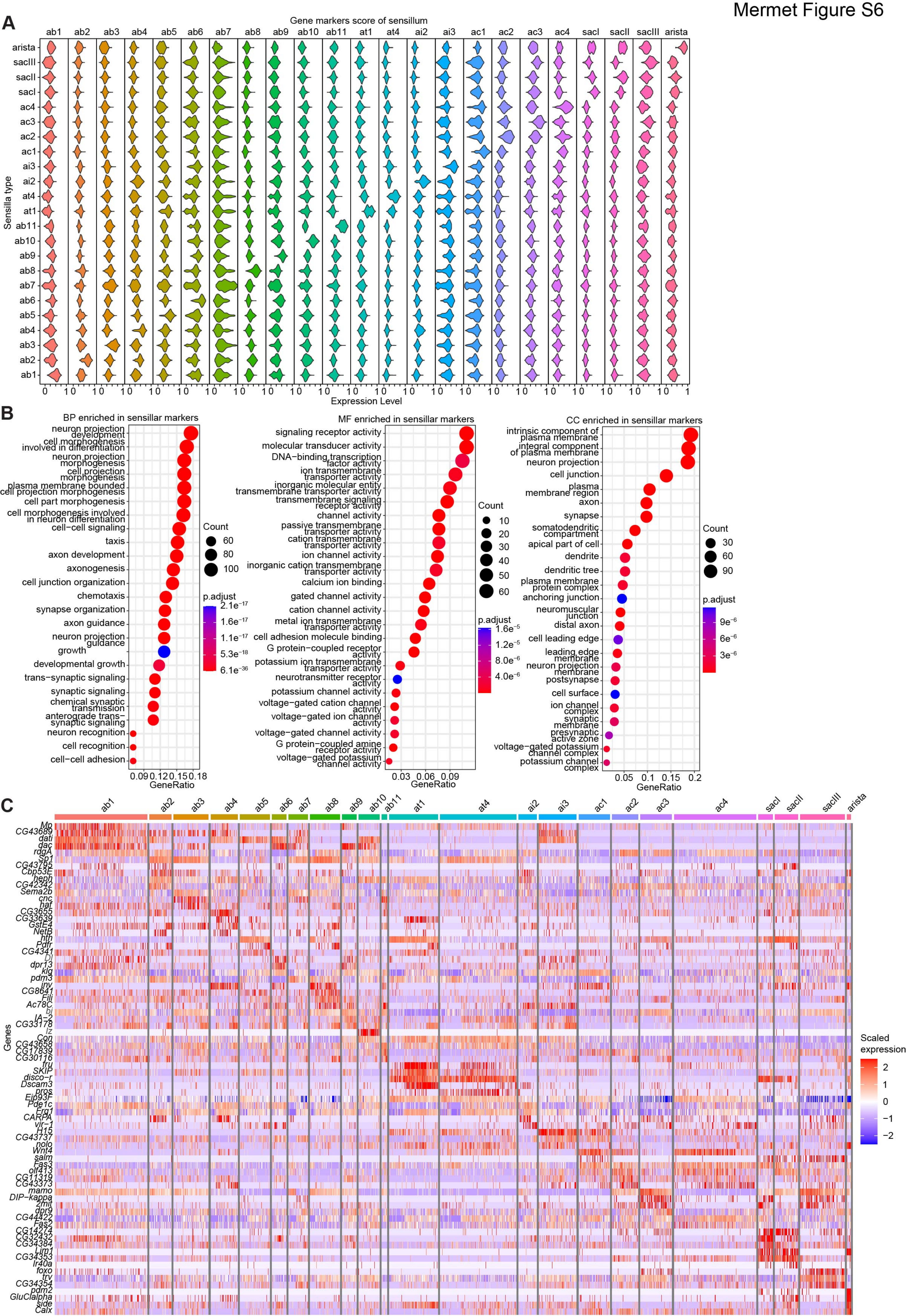

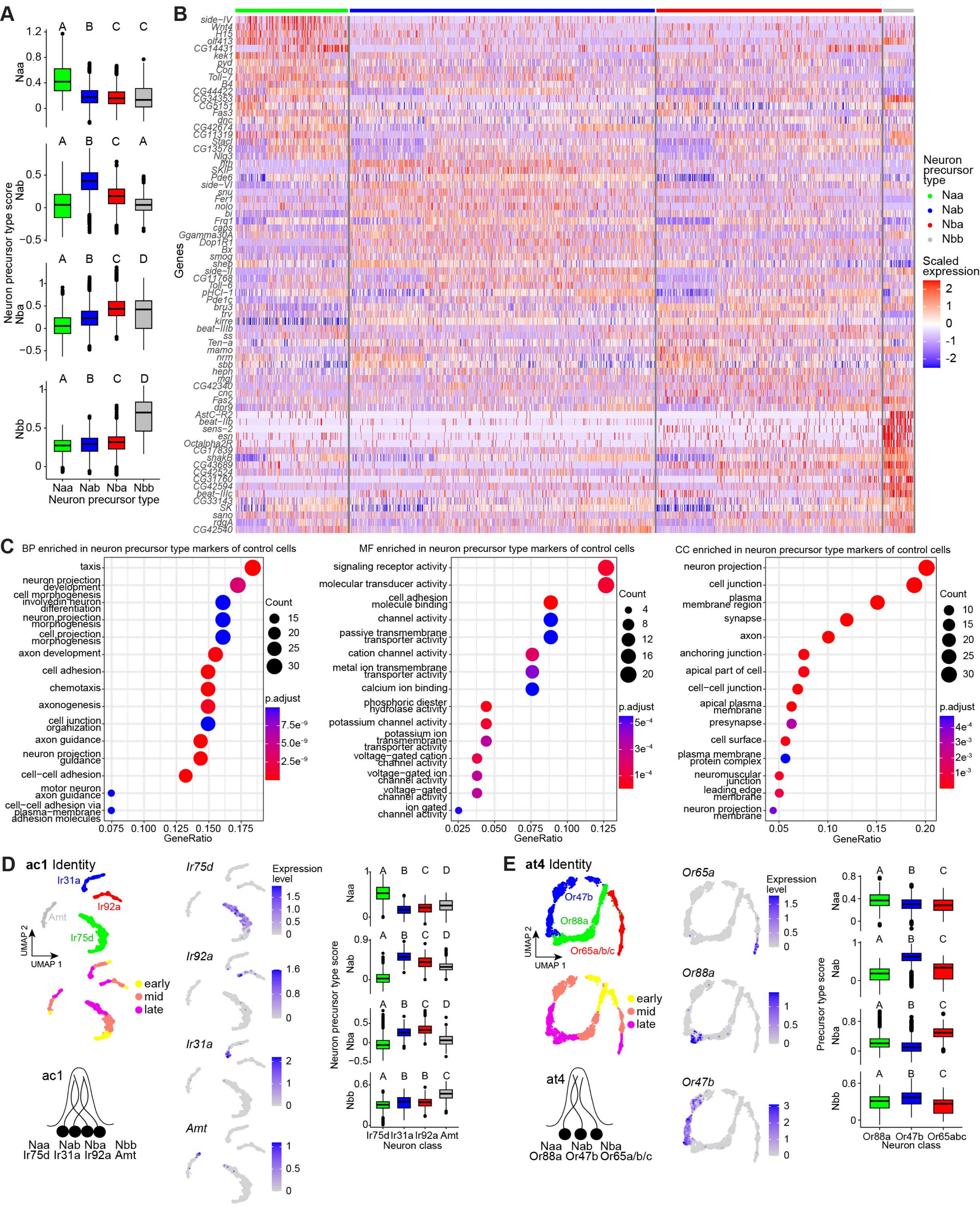
A
B
C
D
E
Neuron precursor type score
Naa
Nab
Nba
Nbb
Neuron precursor type
Genes
Neuron precursor type
Scaled expression
BP enriched in neuron precursor type markers of control cells
MF enriched in neuron precursor type markers of control cells
CC enriched in neuron precursor type markers of control cells
GeneRatio
Count
p.adjust
ac1 Identity
Ir75d
Ir92a
Ir31a
Amt
Expression level
UMAP 1
UMAP 2
early
mid
late
Neuron class
at4 Identity
Or65a
Or88a
Or47b
Or65a/b/c
Precursor type score

Mermet Figure S8

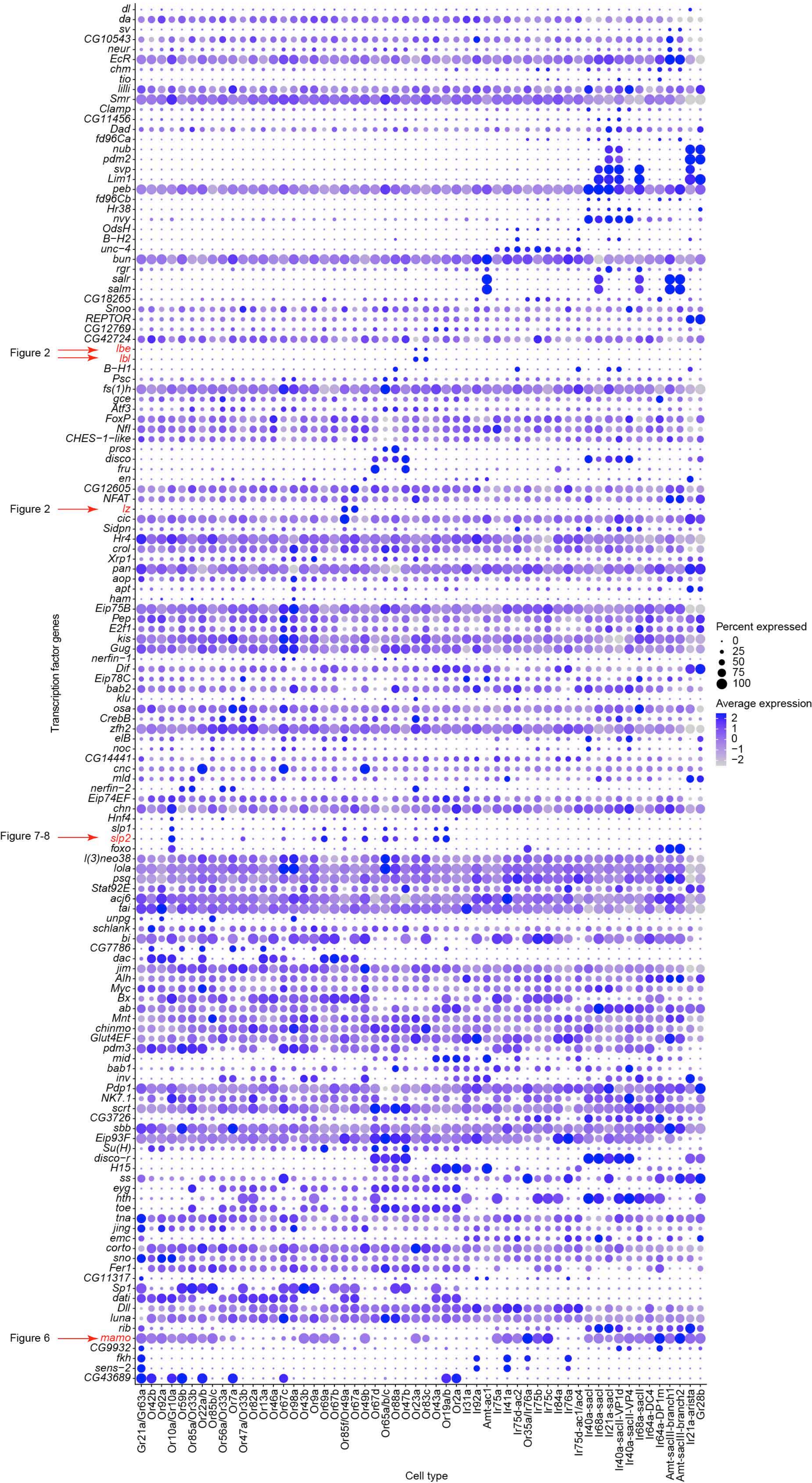

### Mermet Figure S9

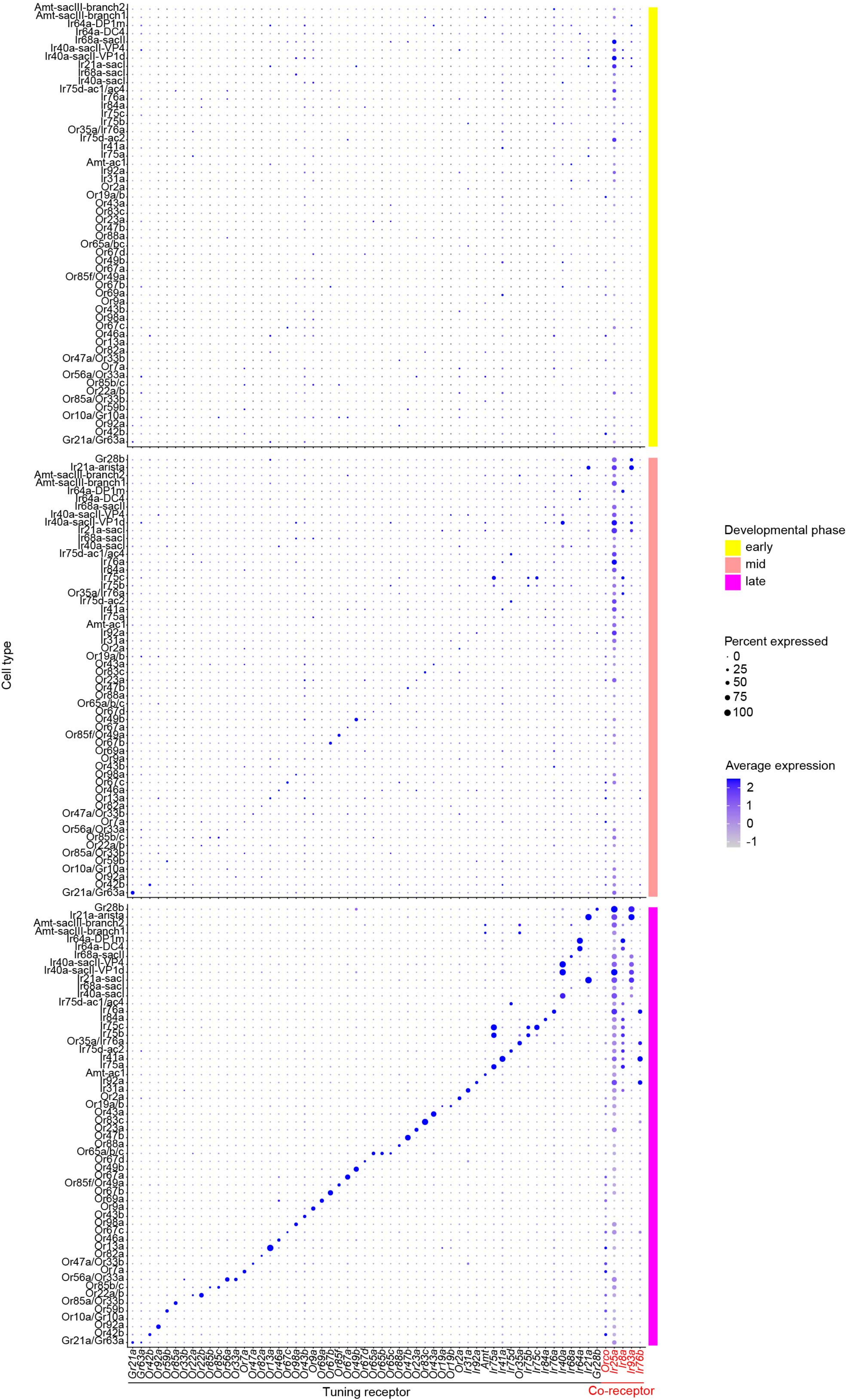

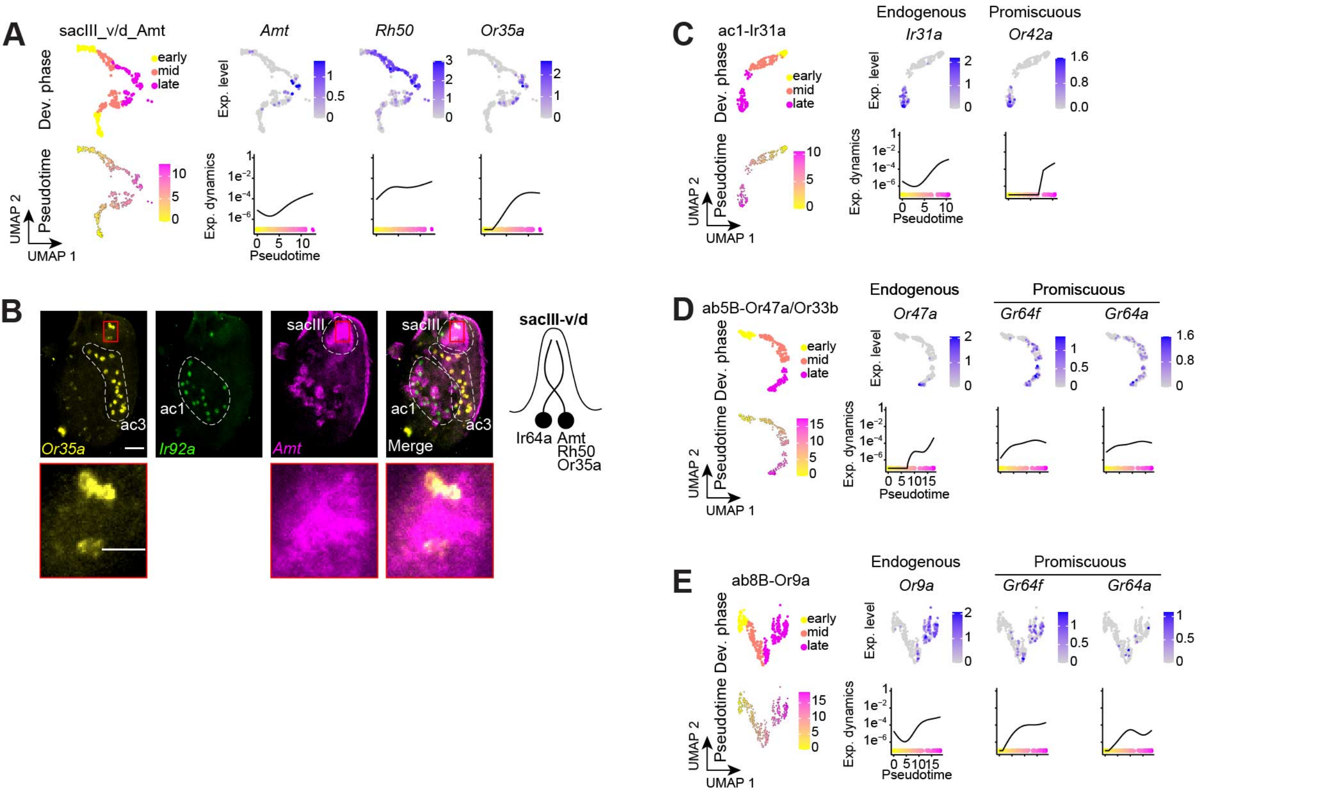
A
sacIII_v/d_Amt
Dev. phase
early
mid
late
Pseudotime
UMAP 2
UMAP 1
Amt
Rh50
Or35a
Exp. level
Exp. dynamics
Pseudotime
B
Or35a
Ir92a
Amt
Merge
ac1
ac3
sacIII
sacIII-v/d
Ir64a
Amt
Rh50
Or35a
C
ac1-Ir31a
Endogenous
Ir31a
Promiscuous
Or42a
D
ab5B-Or47a/Or33b
Endogenous
Or47a
Promiscuous
Gr64f
Gr64a
E
ab8B-Or9a
Endogenous
Or9a
Promiscuous
Gr64f
Gr64a

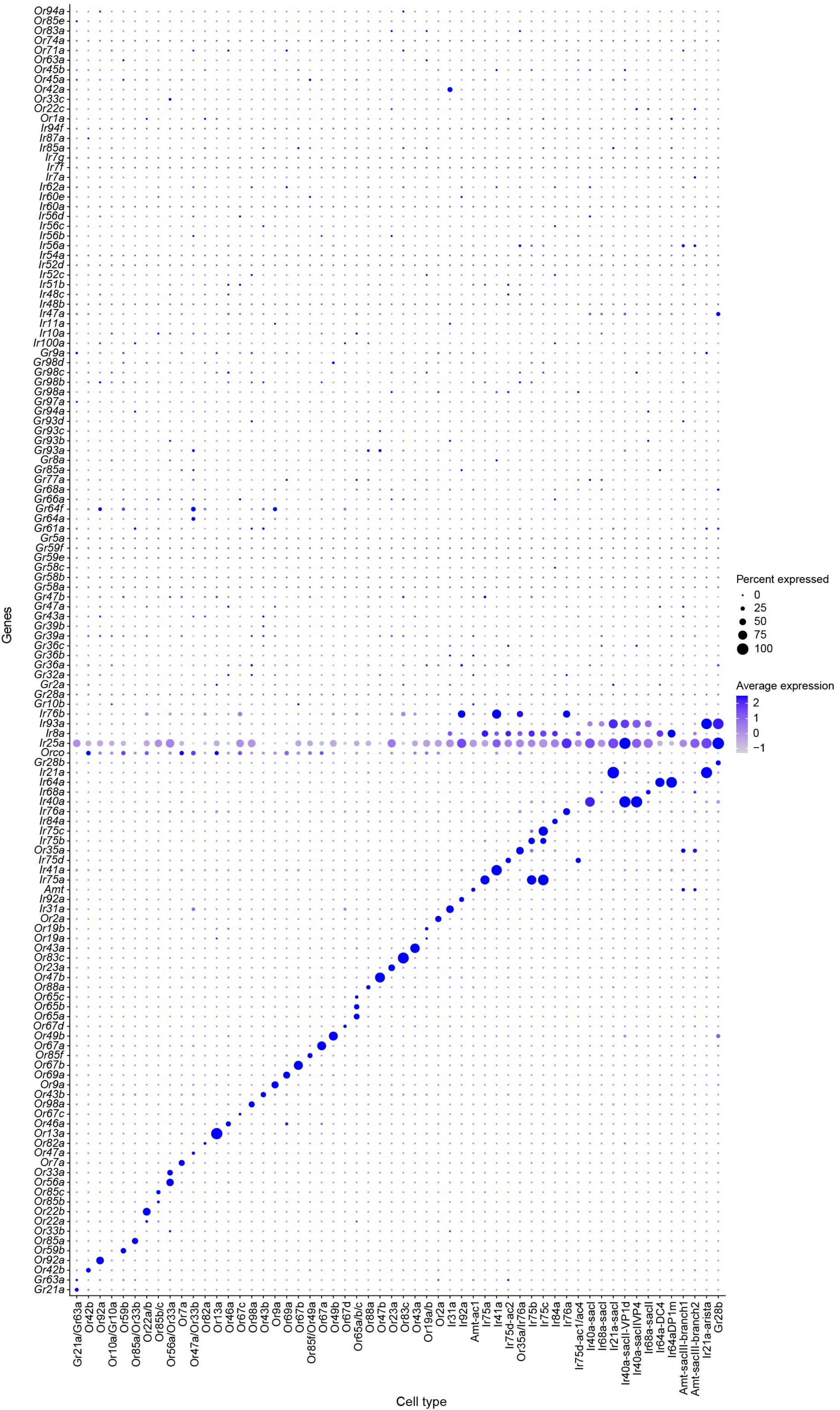
Genes
Cell type
Percent expressed
0
25
50
75
100
Average expression
2
1
0
−1
Or94a
Or85e
Or83a
Or74a
Or71a
Or63a
Or45b
Or45a
Or42a
Or33c
Or22c
Or1a
Ir94f
Ir87a
Ir85a
Ir7g
Ir7f
Ir7a
Ir62a
Ir60e
Ir60a
Ir56d
Ir56c
Ir56b
Ir56a
Ir54a
Ir52d
Ir52c
Ir51b
Ir48c
Ir48b
Ir47a
Ir11a
Ir10a
Ir100a
Gr9a
Gr98d
Gr98c
Gr98b
Gr98a
Gr97a
Gr94a
Gr93d
Gr93c
Gr93b
Gr93a
Gr8a
Gr85a
Gr77a
Gr68a
Gr66a
Gr64f
Gr64a
Gr61a
Gr5a
Gr59f
Gr59e
Gr58c
Gr58b
Gr58a
Gr47b
Gr47a
Gr43a
Gr39b
Gr39a
Gr36c
Gr36b
Gr36a
Gr32a
Gr2a
Gr28a
Gr10b
Ir76b
Ir93a
Ir8a
Ir25a
Orco
Gr28b
Ir21a
Ir64a
Ir68a
Ir40a
Ir76a
Ir84a
Ir75c
Ir75b
Or35a
Ir75d
Ir41a
Ir75a
Amt
Ir92a
Ir31a
Or2a
Or19b
Or19a
Or43a
Or83c
Or23a
Or47b
Or88a
Or65c
Or65b
Or65a
Or67d
Or49b
Or67a
Or85f
Or67b
Or69a
Or9a
Or43b
Or98a
Or67c
Or46a
Or13a
Or82a
Or47a
Or7a
Or33a
Or56a
Or85c
Or85b
Or22b
Or22a
Or33b
Or85a
Or59b
Or92a
Or42b
Gr63a
Gr21a
Gr21a/Gr63a
Or42b
Or92a
Or10a/Gr10a
Or59b
Or85a/Or33b
Or22a/b
Or85b/c
Or56a/Or33a
Or7a
Or47a/Or33b
Or82a
Or13a
Or46a
Or67c
Or98a
Or43b
Or9a
Or69a
Or67b
Or85f/Or49a
Or67a
Or49b
Or67d
Or65a/b/c
Or88a
Or47b
Or23a
Or83c
Or43a
Or19a/b
Or2a
Ir31a
Ir92a
Amt-ac1
Ir75a
Ir41a
Ir75d-ac2
Or35a/Ir76a
Ir75b
Ir75c
Ir84a
Ir76a
Ir75d-ac1/ac4
Ir40a-sacI
Ir68a-sacI
Ir21a-sacI
Ir40a-sacII-VP1d
Ir40a-sacIIVP4
Ir68a-sacII
Ir64a-DC4
Ir64aDP1m
Amt-sacIII-branch1
Amt-sacIII-branch2
Ir21a-arista
Gr28b

Mermet Figure S12

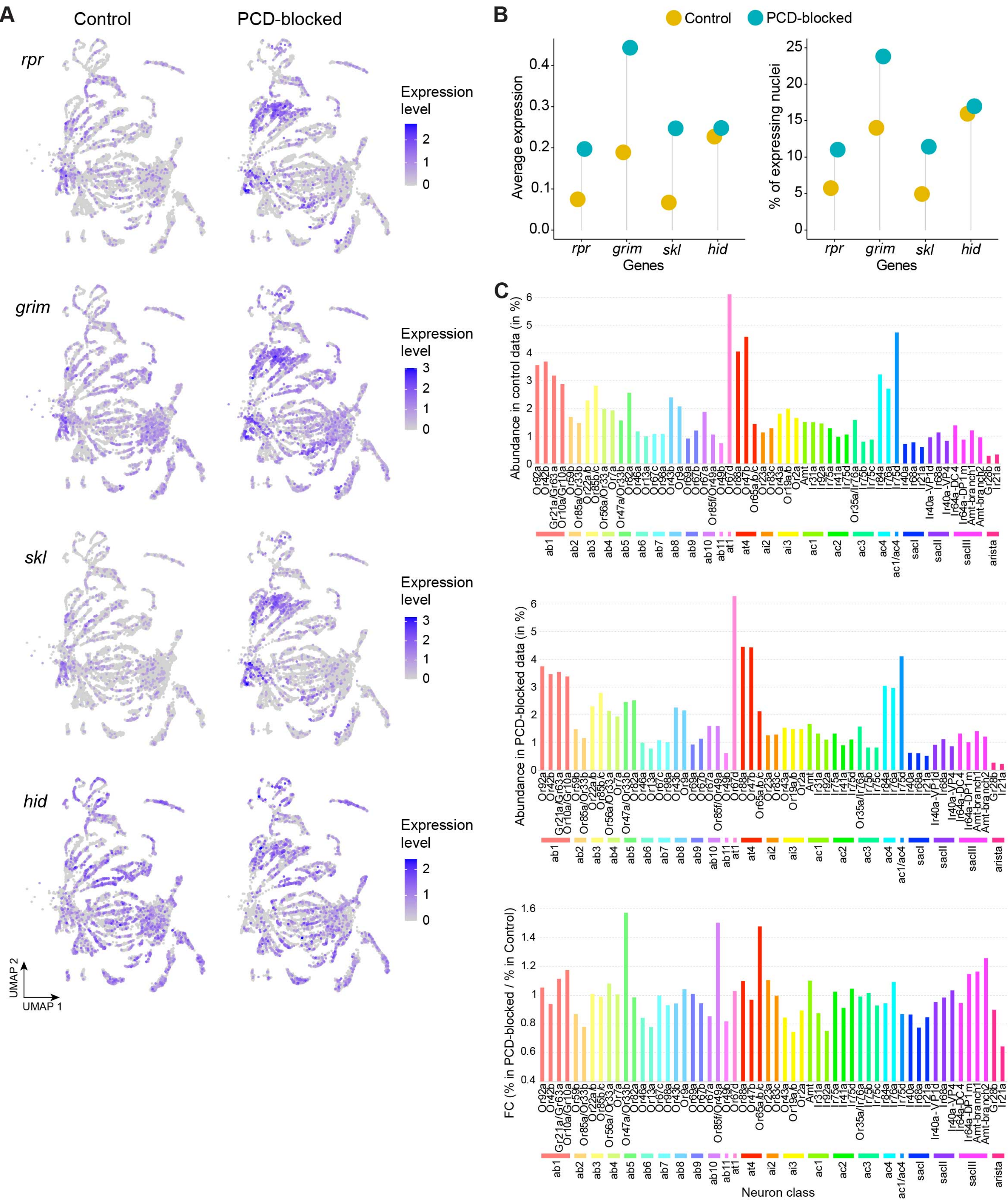

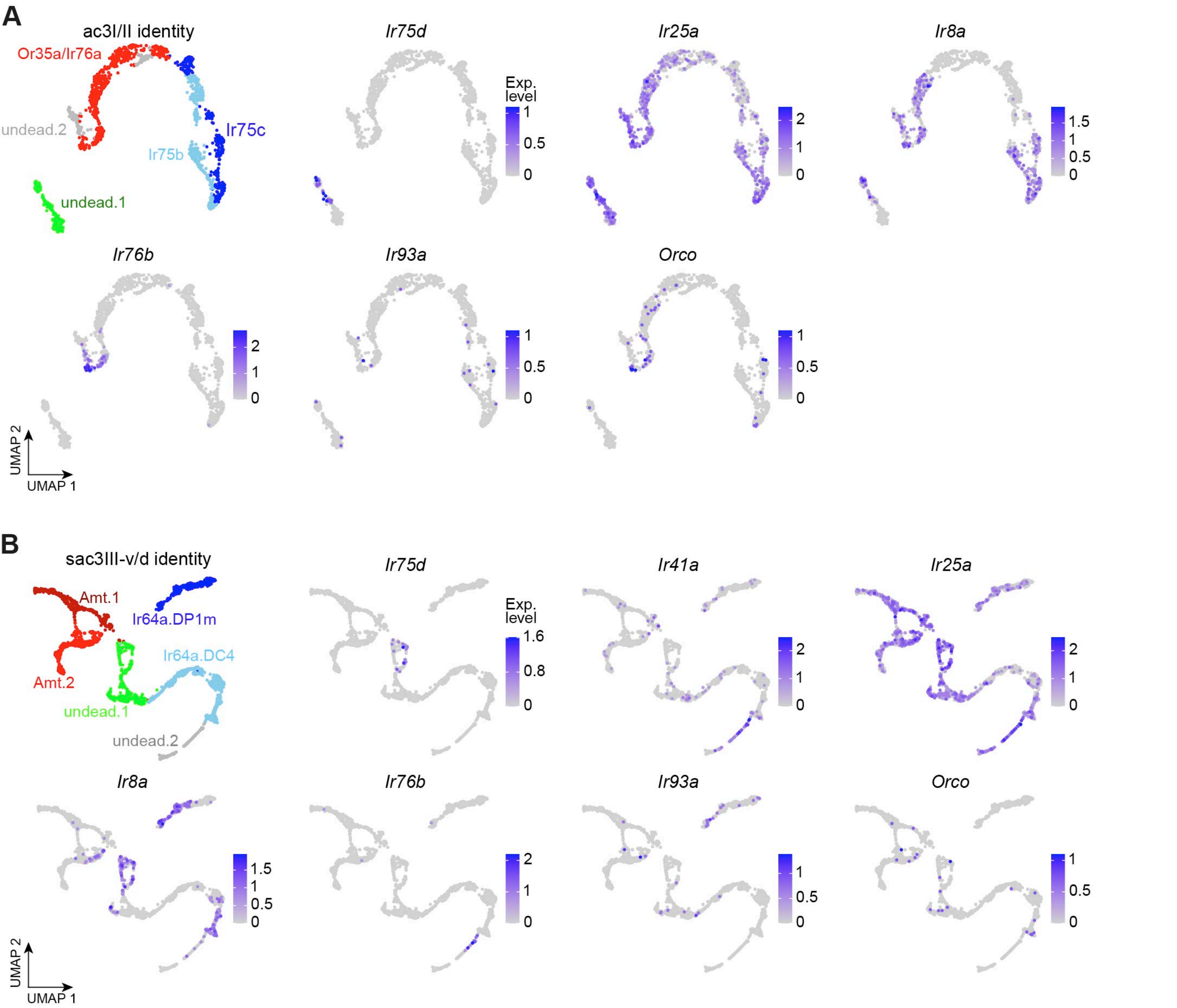
A
ac3I/II identity
Or35a/Ir76a
undead.2
Ir75c
Ir75b
undead.1
Ir75d
Exp. level
1
0.5
0
Ir25a
2
1
0
Ir8a
1.5
1
0.5
0
Ir76b
2
1
0
Ir93a
1
0.5
0
Orco
1
0.5
0
UMAP 2
UMAP 1
B
sac3III-v/d identity
Amt.1
Ir64a.DP1m
Ir64a.DC4
Amt.2
undead.1
undead.2
Ir75d
Exp. level
1.6
0.8
0
Ir41a
2
1
0
Ir25a
2
1
0
Ir8a
1.5
1
0.5
0
Ir76b
2
1
0
Ir93a
1
0.5
0
Orco
1
0.5
0
UMAP 2
UMAP 1

Mermet Figure S14

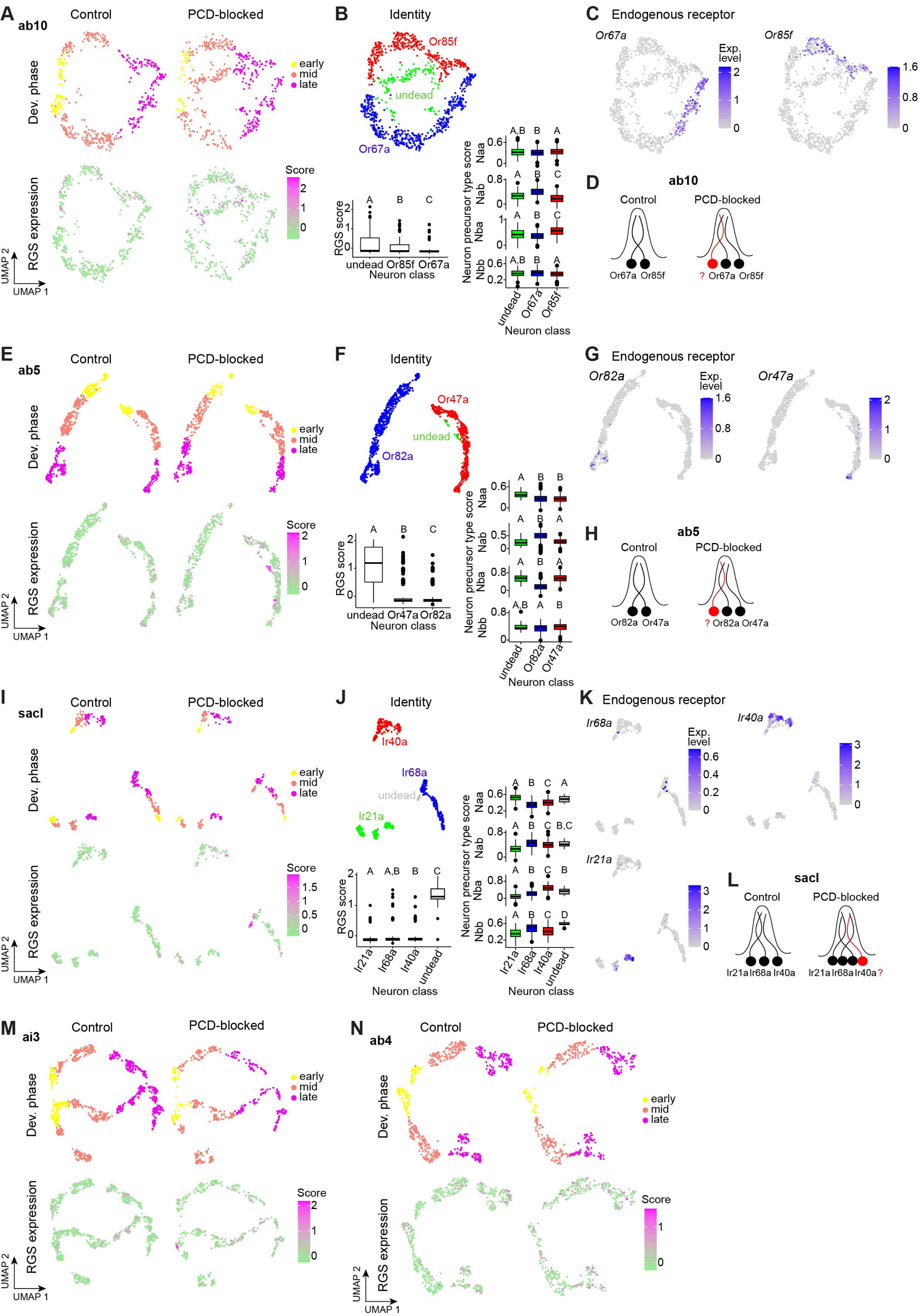

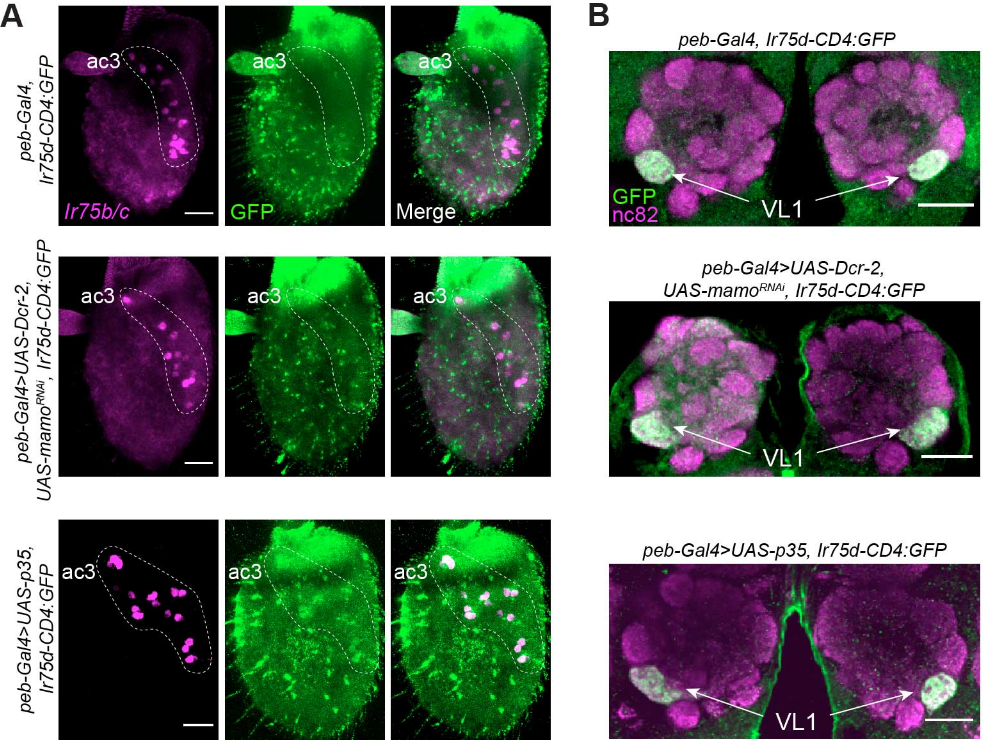
A
peb-Gal4, Ir75d-CD4:GFP
ac3
Ir75b/c
GFP
Merge
peb-Gal4>UAS-Dcr-2, UAS-mamo^RNAi, Ir75d-CD4:GFP
peb-Gal4>UAS-p35, Ir75d-CD4:GFP
B
peb-Gal4, Ir75d-CD4:GFP
GFP
nc82
VL1
peb-Gal4>UAS-Dcr-2, UAS-mamo^RNAi, Ir75d-CD4:GFP
peb-Gal4>UAS-p35, Ir75d-CD4:GFP
